# Balancing selection via life-history trade-offs maintains an inversion polymorphism in a seaweed fly

**DOI:** 10.1101/648584

**Authors:** Claire Mérot, Violaine Llaurens, Eric Normandeau, Louis Bernatchez, Maren Wellenreuther

**Affiliations:** Département de biologie, Institut de Biologie Intégrative et des Systèmes (IBIS), Université Laval, Canada; ISYEB (UMR 7205 CNRS/MNHN/SU/EPHE), Museum National d’Histoire Naturelle, Paris, France; The New Zealand Institute for Plant & Food Research Ltd, Nelson, New Zealand; School of Biological Sciences, University of Auckland, Auckland, New Zealand

**Keywords:** chromosomal inversion, experimental evolution, Individual-based model, sexual selection -structural variation

## Abstract

How genetic diversity is maintained in natural populations is an evolutionary puzzle. Over time, genetic variation within species can be eroded by drift and directional selection, leading to the fixation or elimination of alleles. However, some loci show persistent variants at intermediate frequencies for long evolutionary time-scales, implicating a role of balancing selection, but studies are seldom set up to uncover the underlying processes. Here, we identify and quantify the selective pressures involved in the widespread maintenance of an inversion polymorphism in the seaweed fly *Coelopa frigida*, using an experimental evolution approach to estimate fitness associated with different allelic combinations. By precisely evaluating reproductive success and survival rates separately, we show that the maintenance of the polymorphism is governed by a life-history trade-off, whereby each inverted haplotype has opposed pleiotropic effects on survival and reproduction. Using numerical simulations, we confirm that this uncovered antagonism between natural and sexual selection can maintain inversion variation in natural populations of *C. frigida*. Moreover, our experimental data highlights that inversion-associated fitness is affected differently by sex, dominance and environmental heterogeneity. The interaction between these factors promotes polymorphism maintenance through antagonistic pleiotropy. Taken together, our findings indicate that combinations of natural and sexual selective mechanisms enable the persistence of diverse trait in nature. The joint dynamics of life history trade-offs and antagonistic pleiotropy documented here is likely to apply to other species where large phenotypic variation is controlled by structural variants.

**Significance statement:** Persistence of chromosomal rearrangements is widespread in nature and often associated with divergent life-history traits. Understanding how contrasted life-history strategies are maintained in wild populations has implications for food production, health and biodiversity in a changing environment. Using the seaweed fly *Coelopa frigida,* we show that a polymorphic chromosomal inversion is maintained by a trade-off between survival and reproduction, and thus provide empirical support for a role of balancing selection *via* antagonistic pleiotropy. This mechanism has long been overlooked because it was thought to only apply to a narrow range of ecological scenarios. These findings empirically reinforce the recent theoretical predictions that co-interacting factors (dominance, environment and sex) can lead to polymorphism maintenance by antagonistic pleiotropy and favour life-history variation.

## Introduction

The selective mechanisms involved in the maintenance of long-term polymorphism in the face of genetic drift often remain unknown. Early assessments of heritable diversity, as well as recent empirical genomic and theoretical studies, have all emphasised the importance of balancing selection in promoting within-species diversity and maintaining several alleles at intermediate frequencies within a species (1–5). The best-documented balancing selection regimes are overdominance, where heterozygotes benefit from a higher survival compared to homozygotes, and negative frequency-dependent selection, where allelic fitness decreases with increasing frequency, resulting in both cases in the protection of rare alleles. Balancing selection may also arise from the combination of opposing selection pressures, each favouring different alleles at polymorphic loci (6), for example via spatially or temporally-varying selection, where the fitness of the different alleles varies among heterogeneous habitats or seasons (7–11), or sexually-antagonistic selection, where allelic fitness varies between sexes (12–15).

However, the mechanisms underlying balanced polymorphisms in the wild are still largely unidentified, partly because the fitness associated with different genotypes is affected by several interacting life history factors. Individual fitness results from complex trait combinations of survival, longevity, reproductive success and fecundity, which can be under opposed selective pressures. For example, one allele can increase survival but weaken reproductive success, while the alternative allele can confer high fertility at the cost of decreasing survival, creating a life history trade-off (16, 17). Such antagonistic pleiotropy increases genetic variation via the maintenance of intra-locus polymorphism (18, 19). Antagonistic pleiotropy has long been considered as a minor contributor to balancing selection because theoretical studies predicted it enables persistent polymorphism only for a limited range of parameters (20–22). Nevertheless, recent models suggest that the role of antagonistic pleiotropy has been underestimated (23–26). Indeed, these new models show that several factors, realistic in natural populations, can promote polymorphism persistence. These factors include trait-specific dominance (i.e. when the level of dominance varies between fitness components (26, 27)), sex-specific selection (i.e. selection strength on each fitness component differs between sexes (23)) and spatially and temporally varying selection (24, 25).

These recent theoretical developments urge for a re-examination of the mechanisms allowing polymorphism maintenance in natural systems and to disentangle their effects on the different components of fitness. For example, detailed work on phenotypic variation in horn size in Soay sheep (*Ovis aries*) showed that antagonistic pleiotropy due to a life-history trade-off between survival and reproductive success at a single locus maintains polymorphism by causing an overall net heterozygote advantage (17). Interestingly, horn size was also under sex-specific selection and involved trait-specific dominance, two factors predicted to contribute to the maintenance of genetic variation. Similarly, in the yellow monkeyflower (*Mimulus guttatus*) variability in flower size is related to antagonistic pleiotropy due to a trade-off between viability and fecundity, with the persistence of the polymorphism being further enhanced by spatial and temporal environmental heterogeneity (24, 28, 29). However, direct empirical evidence of antagonistic pleiotropy enabling long-term polymorphism remains scarce, in part because the different components of fitness are rarely estimated separately.

Here, we focused on the seaweed fly *Coelopa frigida*, whose natural populations are all polymorphic at a large chromosomal inversion on chromosome I (30–32). Recombination within large inversions is supressed between the different rearrangements. Consequently, inversions typically behave as a single locus, with alleles corresponding to the different haplotypic rearrangements (33–35). In *C. frigida,* the two alleles, referred to as *α* and *β*, differ by more than 2.5% sequence divergence in coding regions and are observed at intermediate frequencies in both European and North American *C. frigida* populations, suggesting that the haplotypic rearrangements diverged a long time ago and were maintained ever since by balancing selection (30–32). The widespread excess of *αβ* heterozygotes and the higher egg-to-adult survival of heterozygotes compared to *αα* and *ββ* homozygotes implicates that this polymorphism is partly maintained by overdominance, possibly due to deleterious alleles captured by the inversion (36, 37). Moreover, inversion frequencies correlate with environmental factors such as temperature and local substrate characteristics, suggesting that spatially-varying selection combined with migration also contributes to maintain this polymorphism (30, 31, 38). Furthermore, antagonistic pleiotropy on different fitness components may also favour polymorphism given that the inversion affects adult size (Fig. 1A) and that body size is linked to fertility (39, 40), and development time (41) which could modulate egg-to-adult survival. While the phenotypic effect on females is moderate, *αα* males can be up to three times larger than *ββ* males, but *αα* males can also take up twice as long to develop than *ββ* males (30, 41). This pattern strongly suggests a trade-off between adult size and egg-to-adult development, which may result in a trade-off between fertility and survival. These findings make the inversion polymorphism in *C. frigida* a relevant empirical case to investigate the role of antagonistic pleiotropy in promoting polymorphism and to specifically test interactions with other mechanisms favouring the maintenance of variation.

**Figure 1:**
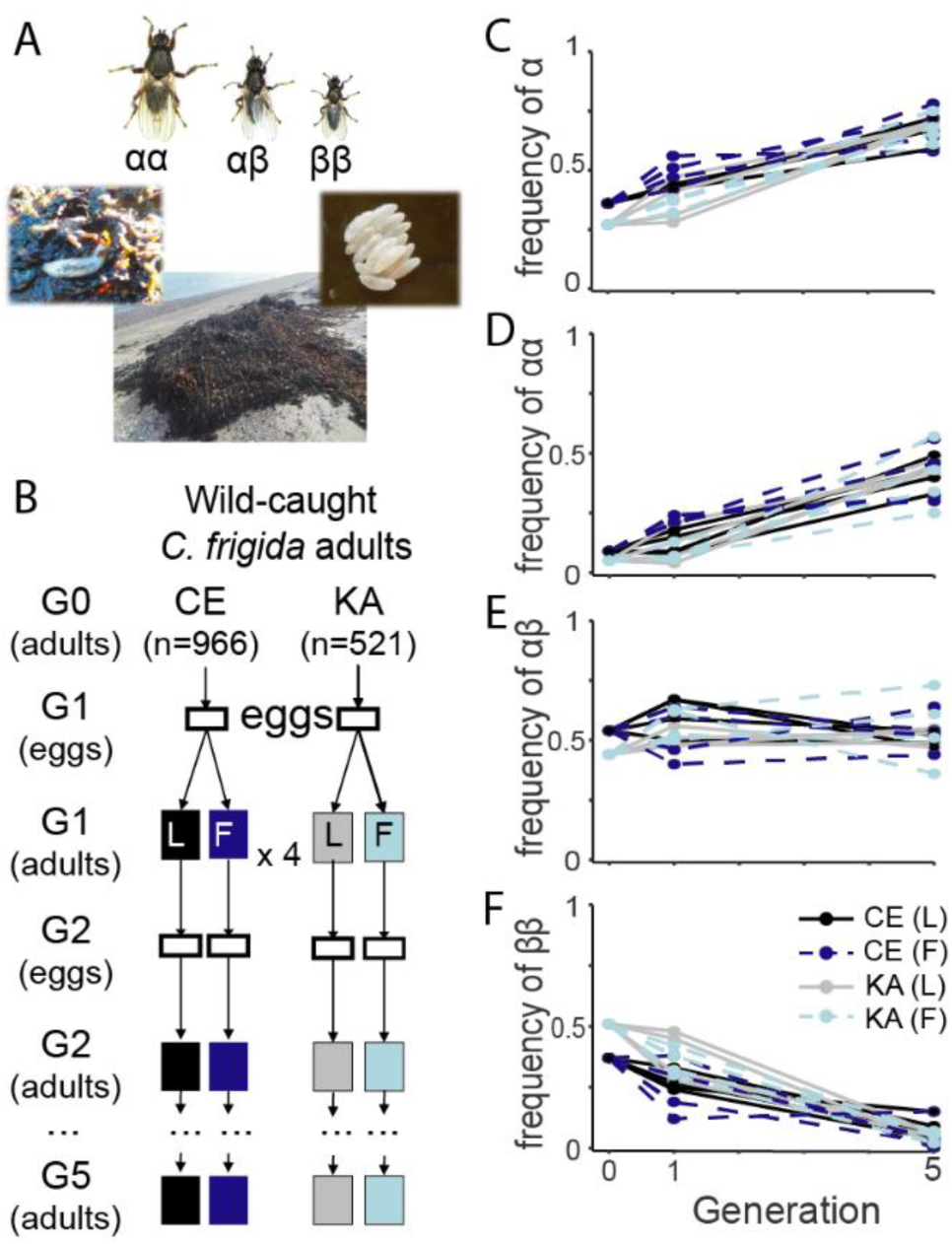
*In vivo* experimental evolution of *Coelopa frigida* and inversion dynamics. ***(A)** C. frigida* is a seaweed fly inhabiting seaweed wrackbeds that are accumulating and decomposing on the shoreline. Larva are exclusively restricted to this wrackbed substrate and adults are generally found crawling on or within the decomposing seaweed on which they lay eggs in clusters, although they can at times stray away from the wrackbed. Size variation in adult males is associated with the three genotypes of the inversion. ***(B)*** Overview of the laboratory evolution experiment design. Starting with wild populations collected from two locations (CE & KA) in Québec (Canada), we raised 16 replicated experimental populations separately over 5 generations (denoted as G), either on a substrate dominated by Laminariaceae (*L*) or Fucaceae (*F*). Eggs and adults were genotyped for an SNP marker associated with the inversion to infer genotype frequencies. ***(C-F)*** The frequency of the inversion allele *α* increased between generation 0, 1 and 5 as *ββ* homozygotes were replaced by *αα* homozygotes. The same trend was observed in all 16 replicates for both KA and CE origins and on both substrates.

We combine experimental evolution and simulations to investigate the mechanisms of balancing selection underlying the maintenance of the inversion polymorphism in *Coelopa frigida* and to decipher the role of antagonistic pleiotropy in this process by estimating the effect of inversion alleles on different fitness components. First, we use experimental evolution (Fig. 1B) to follow the inversion frequencies for 5 generations and to estimate egg-to-adult survival and reproductive success. Second, realistic numerical simulations based on these experimentally-estimated parameters (Fig. 3A) allow quantifying the contribution of antagonistic pleiotropy in the maintenance of this inversion polymorphism. Both the experiments and the simulations support the hypothesis that a life-history trade-off mediates balancing selection maintaining the inversion polymorphism, and that this trade-off is also affected by other factors, namely sex-specific selection and trait-specific dominance. Finally, we expand our simulation model to characterize how the combination of different selective mechanisms favours the persistence of this polymorphism *via* antagonistic pleiotropy.

## Results

### Inversion dynamics during *in vivo* experimental evolution

During the experiment, the frequency of the α allele increased significantly from 27-36% (initial frequency in natural populations) to 58-75% (fifth generation, Fig. 1C, Table S1-2). This pattern was observed in all 16 replicates, with no significant effect of the substrate. Initial differences in frequency between the two origins were lost at generation 5 (Fig. 1C-F, Table S2). The increase in *α* frequency stemmed from a sharp increase of *αα* homozygotes and a reduction in *ββ* homozygotes (Fig. 1D-F). This suggests that the *α* allele confers higher fitness compared to the *β* allele in these experimental conditions, either because of higher survival, higher reproductive success, or both. Proportions of *αβ* heterozygotes remained stable around 50% throughout the experiment (Fig. 1E), suggesting that *αβ* frequencies may be stabilized by overdominance.

To disentangle the fitness components for the three genotypes and explore dominance effects, we genotyped the adults and the eggs at each generation for a subset of four replicates (Table S1) and monitored frequency deviations due to biased survival or non-random reproduction. In combination with follow-up experiments and simulations, we tested for antagonistic pleiotropy associated with the inversion by precisely evaluating four fitness parameters in the three genotypes: egg-to-adult survival, development time, female fecundity and male reproductive success.

### The β allele confers a viability advantage during the larval stage

Egg-to-adult relative survival of *αα* homozygotes was significantly lower than *αβ* heterozygote and *ββ* homozygote survival, and this difference was pronounced (20% on average, Fig. 2A, Table 1). Therefore, the increase of *αα* frequency during the experiments is a paradox given the lower viability of this genotype. Here, *ββ* and *αβ* relative survival rate did not differ significantly (mean difference 2%, *p* = 0.99), suggesting a dominance effect of the *β* allele on egg-to-adult survival. This contrasts with the general finding of overdominance in European populations of *C. frigida*, where *αβ* heterozygotic larvae survive better than both homozygotes (36, 37, 42). Yet, the high *αβ* relative survival is known to be enhanced by increased competitive conditions and, therefore, the performance of homozygotes in our data may be explained by the low-density conditions maintained in our experiment. We also calculated relative sex-specific survival rate of the three genotypes (Fig. S1-S2, Table 1). Although these estimates should be interpreted cautiously given their large variance, our data were consistent with previous estimates on *C. frigida* populations raised at low density (37) (Table S3): overdominance of *αβ* was observed for male survival while *ββ* females tended to outperform *αβ* females (Fig. S1, Table S3), suggesting that the effects of the inversion on viability as well as on the dominance relationship differs between sexes.

**Table 1:** Fitness parameters depending on sex (M=male, F-=female) and genotype (*αα, αβ* and *ββ*). Within each cell, the first line is the parameter name, the second line is the numeric values inferred from the experiment and used by default in most simulations exploring a realistic set of parameters, the third line is a conceptualization of the parameter based on Zajitschek & Connallon [21] used for the simulations exploring a theoretical space of parameters, with s_m_, s_f_ being the coefficients of selection for survival in males and females, and t_m_ and t_f_ being the coefficients of selection for reproduction in males and females. *H_s_* is the coefficient of dominance for survival and *H_t_* is the coefficient of dominance for reproduction. Darker cell backgrounds indicate the best value(s) for each parameter and denote sex-dependent and trait-dependent dominance effect.

| Sex | Fitness component | Genotype at the inversion locus |  |  |
| --- | --- | --- | --- | --- |
| | | $\alpha\alpha$ | $\alpha\beta$ | $\beta\beta$ |
| M | Egg-to-adult relative survival | $S_{aa-m}$<br>0.81<br>$1-s_m$ | $S_{a\beta-m}$<br>1.0<br>$1-s_m \cdot H_s$ | $S_{\beta\beta-m}$<br>0.88<br>1 |
| | Relative reproductive success | $T_{aa-m}$<br>1.0<br>1 | $T_{a\beta-m}$<br>(0.55)<br>$1-t_m \cdot H_t$ | $T_{\beta\beta-m}$<br>(0.1)<br>$1-t_m$ |
| F | Egg-to-adult relative survival | $S_{aa-f}$<br>0.71<br>$1-s_f$ | $S_{a\beta-f}$<br>0.90<br>$1-s_f \cdot H_s$ | $S_{\beta\beta-f}$<br>1.0<br>1 |
| | Relative reproductive success | $T_{aa-f}$<br>1.0<br>1 | $T_{a\beta-f}$<br>0.97<br>$1-t_f \cdot H_t$ | $T_{\beta\beta-f}$<br>0.87<br>$1-t_f$ |

**Figure 2:**
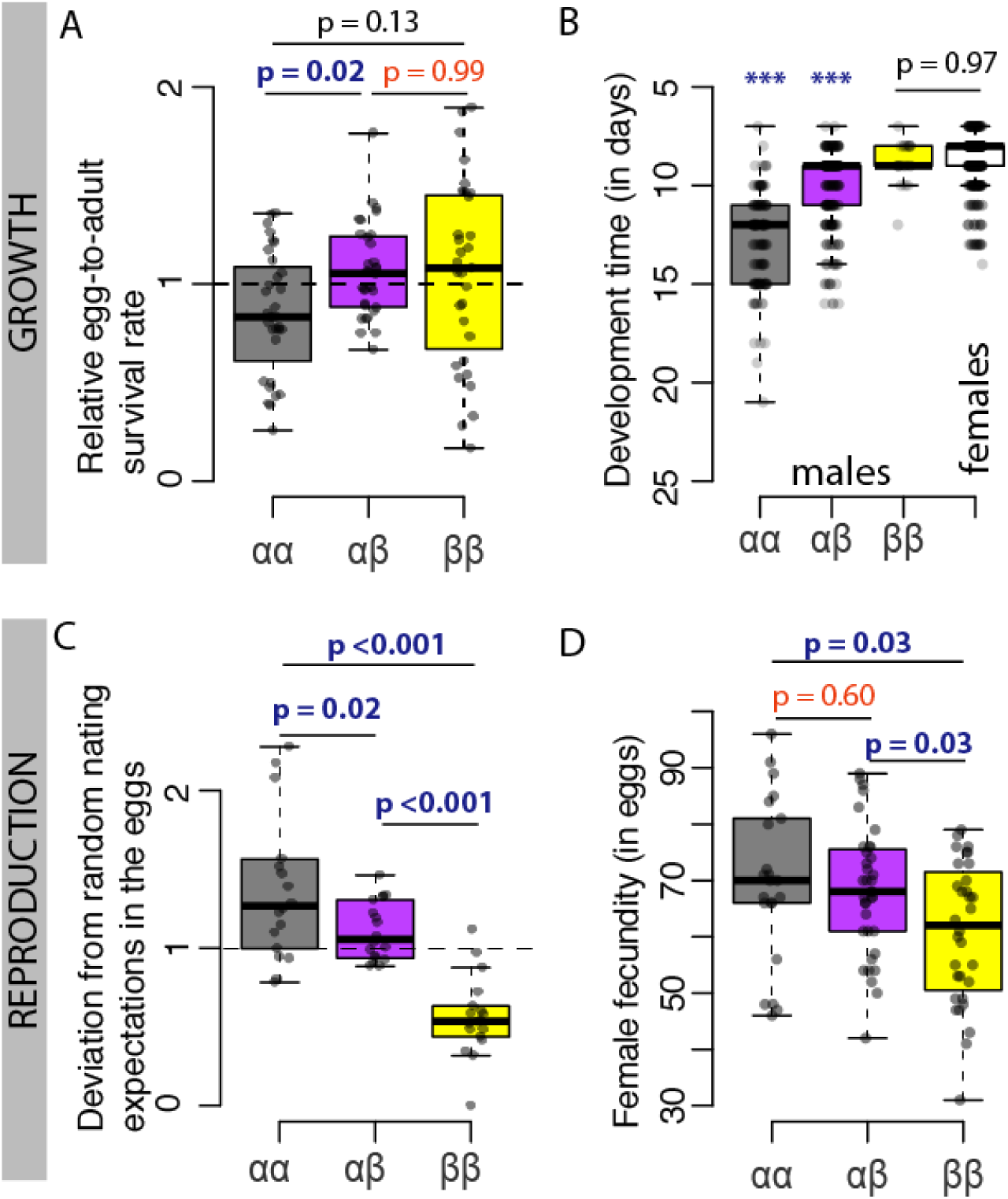
The pleiotropic and antagonistic effects of the inversion on different components of fitness. ***(A)*** Relative egg-to-adult survival rate per genotype, calculated as the deviation of each genotype’s proportion between adults and eggs (males and females were considered together). ***(B)*** Development time, measured as the number of days from the egg to the emerging adult for each combination of genotype and sex (the white box being developmental time for all females given that no significant difference was found between genotypes). ***(C)*** Deviation of genotypic proportions in the eggs relative to the proportions expected under random mating of the previous generation. ***(D)*** Female fecundity, measured as the number of eggs in the first clutch. *P*-values are indicated in blue and represent significant differences in post-hoc pairwise *t*-test. *P*-values indicated in red represent non-significant differences between the heterozygotes and homozygote, suggesting dominance relationships between *α* and *β* alleles.

The inversion also showed a sex-specific effect on the duration of the larval stage, with no significant difference among female genotypes (on average, *αα:* 9.0 days, *αβ:* 8.7 days, *ββ:*9.0 days; *p* = 0.57) but highly significant differences among males (*p* < 0.001, Table S5, Fig. 2B). In our experimental conditions, (*i.e*. 25°C and low-density) the development time was on average 50% longer for *αα* males (12.8 days), and 25% longer for *αβ* males (10.3 days) compared to *ββ* males (8.8 days) or females. Such ordering between genotypes is consistent with previous studies, although absolute values are larger at higher density (41, 43) or lower temperature (pers. obs., CM and MW). Therefore, the *α* allele is expected to confer an additional mortality risk in the natural environment for males by extending larval stage duration and delaying time to the onset of sexual maturity, although this effect is challenging to quantify in the laboratory where the substrate is not limiting.

### The *α* allele confers a reproductive advantage during the adult stage

Egg genotype proportions significantly deviated from proportions expected under random mating, *i.e.* based on the Hardy-Weinberg proportions of the previous generation (combined probabilities, *p* = 0.003). The deviation consistently corresponded to a noticeable excess of *αα* eggs (on average + 43%), and a slight excess of *αβ* eggs (+ 11%) and a corresponding strong deficit of *ββ* eggs (−44%) (Fig. 2C). Those deviations mean that the *α* allele provided an important advantage for reproduction potentially underlying the increase of *α* frequency in our experiment. The excess of *αα* homozygotes in the eggs relatively to expectations under random mating also decreased between generation 1 and 2 and generation 4 or 5, while the proportions of *αα* increased (Fig. S4, Table S6), suggesting a possible negative frequency-dependence effect on reproduction.

The excess of the *α* allele in the eggs was partly explained by a higher fecundity of the females bearing the *α* allele. In a follow-up experiment on 90 females, *αα* and *αβ* females laid significantly more eggs than *ββ* females in their first clutch, with 15% more eggs on average (*αα*: 70 eggs, *αβ*: 68 eggs, *ββ*: 61 eggs -Fig. 2D), possibly due to the larger adult size associated with *α* also observed in females (39). The number of eggs laid by *αβ* females and *αα* females did not differ significantly (Fig. 2D), suggesting that the advantage in female fecundity associated with the *α* allele may be dominant.

The excess of *α* in the eggs may also result from non-random mating favouring *α* males because of their larger body size. Previous experiments in *C. frigida* with two males and one female demonstrated that the largest male sire a disproportionate higher number of the progeny (44). The reasons are two-fold: smaller males lose male-male competition by being dislodged by larger males (45); and, even in single-pair experiments, smaller males are more likely to be rejected by females than larger males, with *ββ* males being 30 % and 20 % less attractive than *αα* and *αβ* males, respectively (40, 42, 46). To estimate male reproductive success in our experiment, we build an individual-based model simulating the evolution of inversion frequencies *in silico* over 5 generations with all parameters drawn from the experiment except for the male relative reproductive success (Fig. 3A, Table1, Table S7). A scenario with equal male reproductive success across genotypes did not fit our experimental data (Fig. 3B), meaning that the difference in female fecundity could not be sufficient to explain the observed rise in *α* frequency and the excess of *αα* in the eggs. The best model fit was achieved by a 10-fold higher mating success in *αα* compared to *ββ* males (*Tββ-m* = 0.1, Fig. 3B-C, Table S8), with co-dominant, intermediate values for *αβ*. Models involving dominance were explored but generally exhibited a lower model fit (Table S8). Given that male mating success is expected to be linked to male size, a model of co-dominance is more appropriate since *αβ* mean size is intermediate between the two homozygote genotypes (30). Male-male competition effects on relative reproductive success could also account for the decrease of *αα* excess in the eggs at generation 4 and 5 if the higher frequency of large *αα* males translates into increased competition from same-size males and lower individual relative success. Adding a negative frequency-dependent mating success associated with *α* to the model provided a good fit to our experimental data but this was not significantly better compared to simpler models (Table S8).

**Figure 3:**
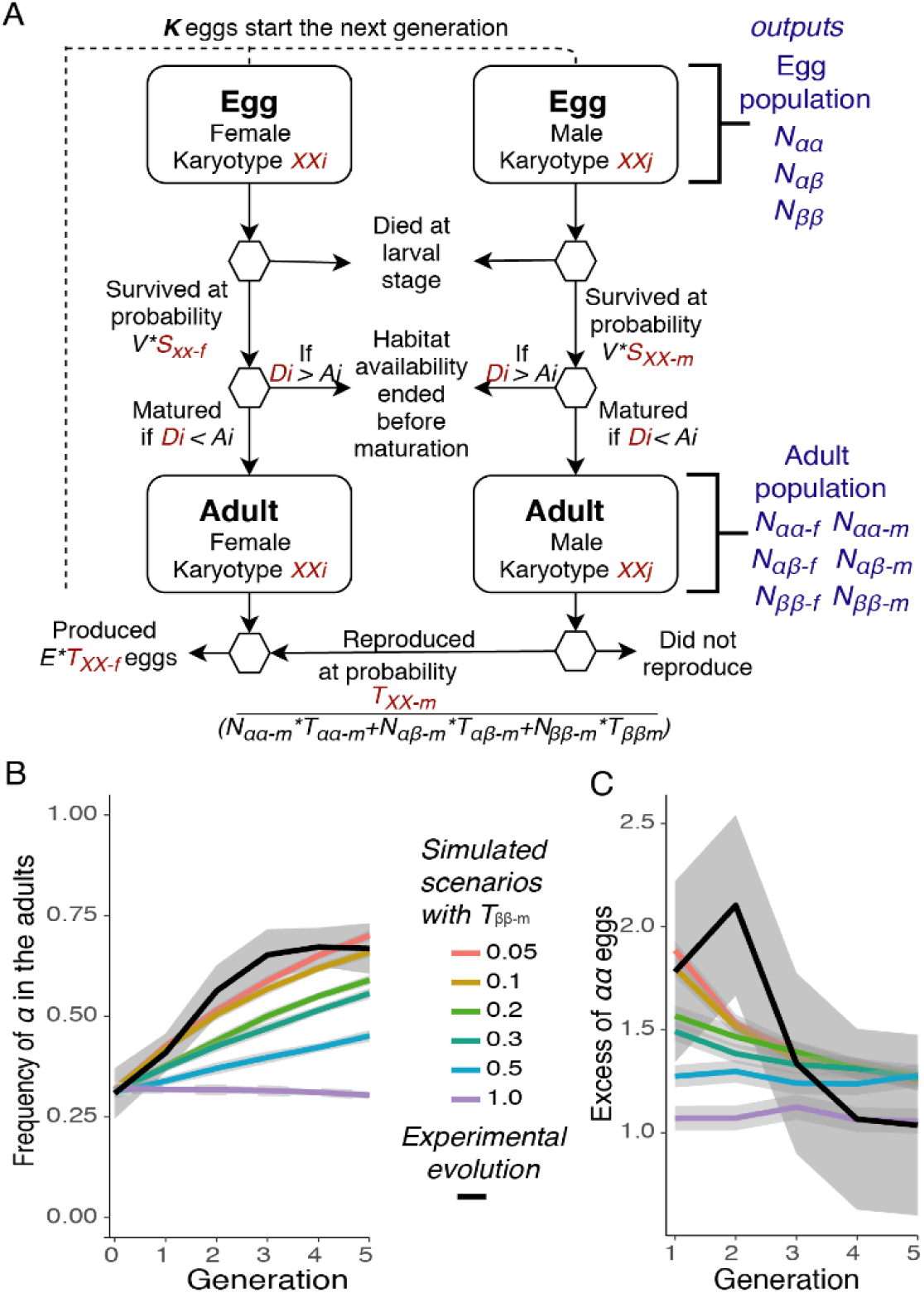
*In silico* evolution of *Coelopa frigida* and inversion dynamics over 5 generations. ***(A)*** Overview of the individual-based simulation model. Each egg is characterized by a genotype and a sex. Eggs survive through the larval stage at a probability determined by the product of *V* (global viability = 0.3) and *S,* the relative survival rate for each combination of sex and genotype. The larva transitions into an adult only if its development time, an individual value (*D_i_*) drawn from a distribution determined by its genotype and sex, is shorter than its individual value of habitat availability (*A_i_*), drawn from a uniform distribution of habitat availability characterized by two parameters, mean duration (*A_mean_*) and variability (*A_var_*). For scenarios simulating laboratory experimental conditions, *A_mean_* is set to a large value (30 days). Adults go through a reproduction phase during which all females mate and lay a number of eggs determined by the product of the fertility parameter *E* (70 eggs) and *T,* the relative female reproductive success by genotype. For each female, a random male is drawn from the pool of adult males at a probability T, the relative male reproductive success, determined by their genotype, and based on the relative proportions of males in the population. Male can reproduce several times. The next generation starts with a subset of *K* eggs representing either the experimental procedure or a limited carrying capacity in nature. ***(B-C)*** Comparing the experiment to simulated scenarios of *in silico* evolution over 5 generations based on experimental parameters but varying male relative reproductive success suggests that relative reproductive success for *ββ* males is 10-fold lower (*T_ββ-m_*=0.1) than for *αα* males (*T_αα-m_* fixed at 1) in experimental conditions. *T_αβ-m_* is the average of the homozygotes’ values (co-dominance model).

### Modelling the impact of antagonistic and pleiotropic effects on inversion dynamics in the wild

To explore how a life-history trade-off modulate inversion frequencies dynamics in nature and to test whether they can contribute to the maintenance of the polymorphism over time, we expanded the individual-based model to 200 generations and used a larger population size. Our experiment allowed us to estimate realistic values for the different life-history parameters. Yet, the increase of *α* frequency to 50-75% in the experiment departs sharply from the lower *α* frequencies generally observed in natural populations (30 – 50 % (30, 38)). This means that there were key differences between wild and experimental conditions that the model needed to take into account. We identified three main differences: (i) our experimental boxes were likely less densely crowded than natural wrackbeds and density is known to strongly affect relative egg-to-adult survival (37), (ii) during the experiment, male-male competition was likely stronger than in nature because of restricted space and synchronised reproduction, which may have favoured large *αα* males, (iii) the experimental substrate was unlimited while in nature access to the resource is frequently disrupted by tides or storm-induced waves, which may prevent slow-developing males to reach adulthood (47). We thus varied and explored the effects of the following three parameters on genotype proportions: (i) relative egg-to-adult survival, (ii) male relative reproductive success, and (iii) duration of habitat availability. We then estimated the probability of maintaining polymorphism *vs.* fixing one allele within both a realistic and theoretical (less constrained) parameter space.

The effect of density on egg-to-adult survival (i) could not account for frequency differences between the wild and the experiment (Fig. 4A). Yet, higher densities shifted the equilibrium proportions towards an excess of heterozygotes as observed many natural populations (4A-D).

**Figure 4:**
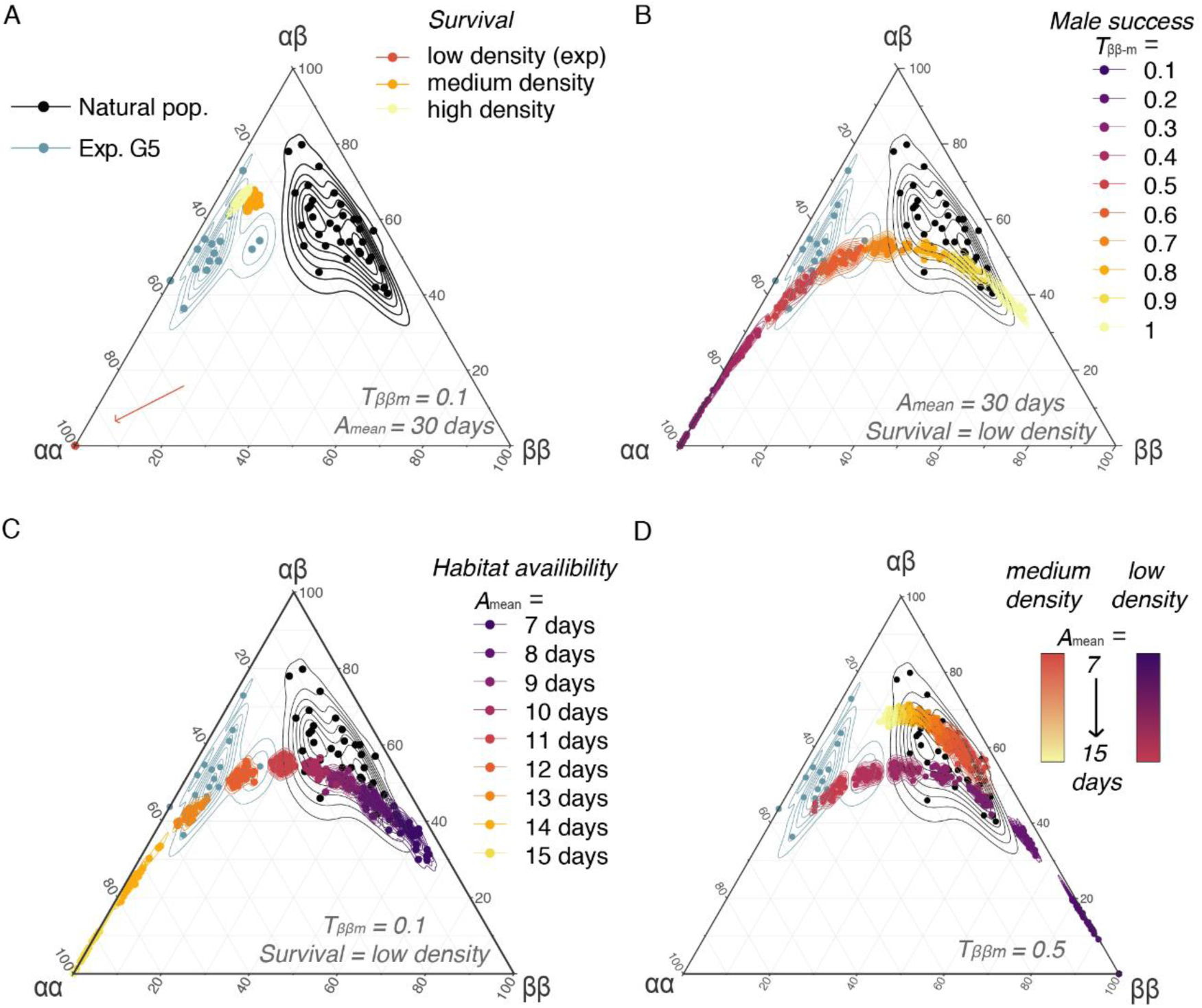
*In silico* evolution of frequencies of inversion genotypes assuming different scenarios exploring the parameter space representative of the conditions encountered by *Coelopa frigida* in the wild. Ternary plots comparing the proportions of the three genotypes in natural populations [29,30], after the 5^th^ generation of our laboratory experiment and at the equilibrium after 200 generations of simulations in scenarios varying ***(A)*** the effect of density, and the related relative survival rate, ***(B)*** the range of values for male relative reproductive success (*T_αα-m_* = 1, *T_αβ-m_* = 1/2*(*T_αα-m_*+ *T_ββ-m_*), *T_ββ-m_*=[0.1-1.0]), ***(C)*** the effect of a limited duration of the habitat availability (*A_mean_* = [7-15 days], *A_var_* = 2 days), and ***(D)*** the combined effects of density and environment for moderate differences of male relative reproductive success (*ββ* male reproductive success *T_ββ-m_*=0.5, *i.e.* two-fold lower than *αα* male reproductive success).

Simulations reducing the relative reproductive success of *αα* males (ii) to about 2-fold (instead of 10-fold in our experiment) recovered an equilibrium of genotypic proportions close to values observed in the wild (Fig. 4B-D), confirming that the increase of *α* frequency in our experiments could be linked to a large reproductive advantage of males carrying the *α* allele, possibly intensified by male-male competition due to restricted experimental space and synchronised reproduction. In the wild, *ββ* males may also have easier access to females by reaching adulthood earlier than *αα* and *αβ* males. However, equal male reproductive success between the three genotypes led to higher *ββ* proportions than what is generally observed in nature and removing the *α*-female fecundity advantage led to the fixation of *β* allele. This suggests that, to some extent, the reproductive advantage conferred by the *α* allele in both males and females, as revealed by the experiment, is also contributing to the persistence of *α*/*β* polymorphism in the wild.

Finally, the duration of habitat availability (iii) appeared to be a prominent factor affecting genotypic proportions and explaining the difference in genotype frequencies between our experiment and wild populations (Fig.4C). This was mediated by a different balance between survival and reproductive advantage. When the habitat availability was shorter, relative survival of *αα* males, and to a lesser extent of *αβ* males, was reduced in comparison to the faster-developing females (all three genotypes) and *ββ* males (Fig. 5B, Fig. S6). It is noteworthy that variation in the duration of habitat availability tended to cover the natural variability in genotypic proportions (Fig. 4C-D), suggesting that spatio-temporal heterogeneity in habitat availability could explain the variable balance of genotypes observed in nature.

**Figure 5.**
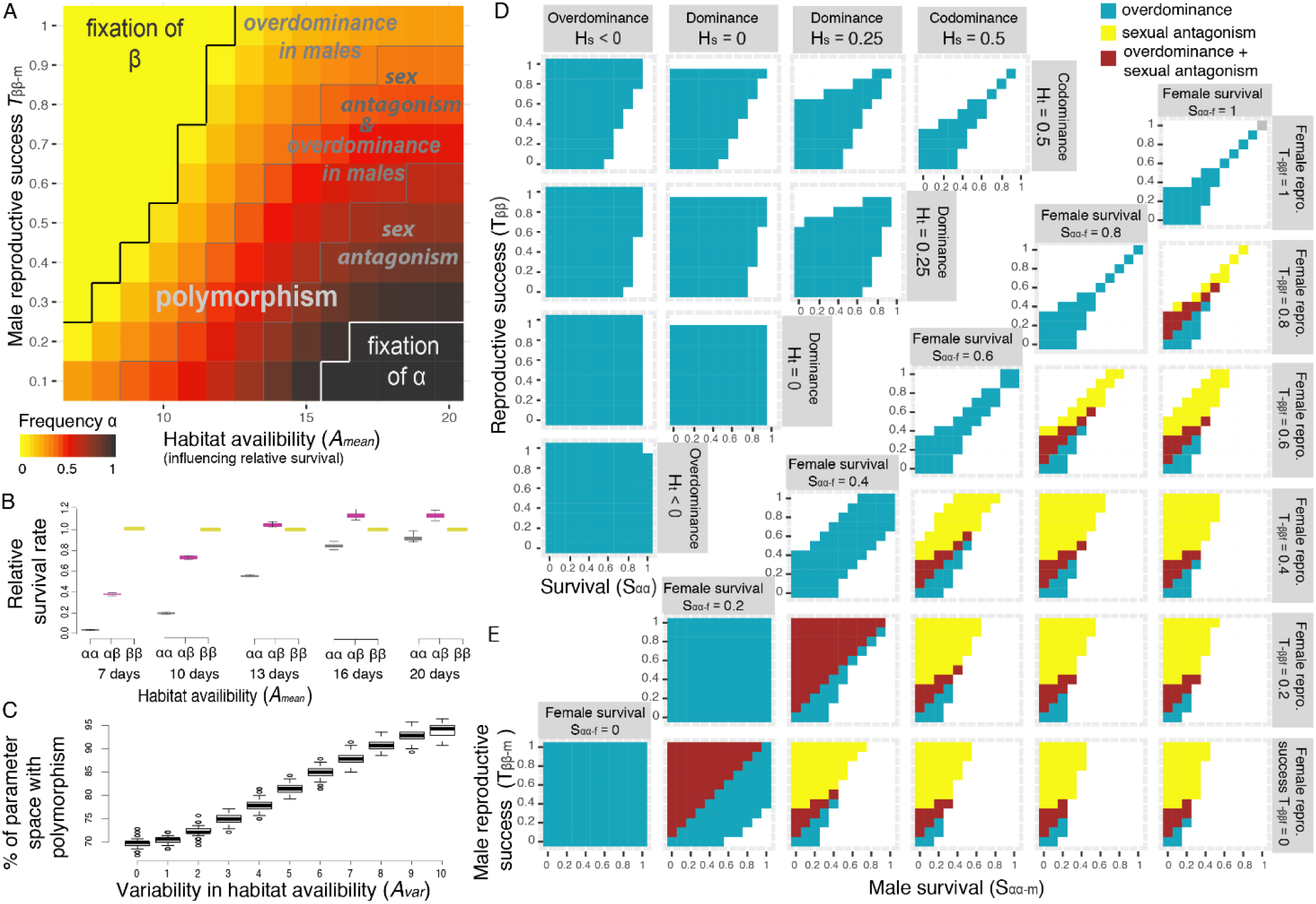
Conditions of maintenance of polymorphism through antagonistic pleiotropy. ***(A)*** Simulations using parameter values motivated by wild populations of *C. frigida* result in intermediate inversion frequencies over a large range of realistic parameters for male relative reproductive success and the duration of habitat availability. Polymorphism is promoted by antagonistic pleiotropy between fitness components which results, at the level of total fitness, in the emergence of overdominance when the heterozygote fitness was higher than both homozygotes’ total fitness, and/or sexual antagonism, when the allele with the highest total fitness was different between sexes. Female reproductive parameter values were based on experimental values, variability in the duration of the habitat (*A_var_*) was set to 2 days and relative survival rates (SXX-s parameters) correspond to low-density conditions. ***(B)*** The duration of habitat availability (*A_mean_*) modulates overall male relative survival by limiting the ability of genotypes with longer development times to reach maturity. Values are normalized relatively to a value of 1 for *ββ* males. ***(C)*** Variability in the duration of habitat availability (*A_var_*) further increases the portion of the parameter space leading to polymorphism persistence. The parameter space if defined by *T_ββ-m_* = [0.1-1.0] and *A_mean_*= [7-20 days], as in panel A. Boxplots show the distribution of 100 replicated simulations. ***(D)*** Simulations within a theoretical parameter space, without sex or an environmental effect, show that overdominance in total fitness emerges as a result of antagonistic pleiotropy, even in the case of co-dominance for each fitness component (top right corner: *H_s_*=0.5, *H_t_*=0.5). Simulations in which dominance for survival (*H_s_*) and reproduction (*H_t_*) are independent show how a reversal of dominance and/or variation in the strength of dominance between components of fitness further expand the range of parameters leading to overdominance and the maintenance of polymorphism. ***(E)*** Outcome of simulations within a theoretical parameter space in which fitness parameters are independent between males and females (*S_αα-f_* is the relative survival of αα females, S*_αα-m_* is the relative survival of αα males, *T_ββ-f_* is the relative reproductive success of ββ females, *T_ββ-m_* is the relative reproductive success of ββ males), without dominance (*H_s_*= *H_t_* = 0.5) or an environmental effect. A sex-specific effect on fitness, even without any sexual antagonism for a given fitness component, expands the range of conditions under which antagonistic pleiotropy favours polymorphism through the emergence of sexual antagonism and sex-specific overdominance at the level of total fitness.

### Polymorphism maintenance through antagonistic pleiotropy

Taken together, the antagonistic pleiotropy among inversion-associated fitness components, as revealed by our experimental evolution approach, can explain the maintenance of the inversion polymorphism in nature, albeit considering a different tuning of the trade-off between survival and reproduction. Simulations exploring these different factors simultaneously showed that the polymorphism is maintained for a wide parameter space of male reproductive success and environmental conditions (Fig. 5A, Fig. S7), even at low density conditions, where heterozygote survival advantage is very weak (Fig. 5B, Fig. S6). Remarkably, when the duration of the habitat availability was more variable, the range of conditions maintaining polymorphism was considerably expanded (Fig. 5C). Also, the frequency of the inversion was buffered around intermediate values (Fig. S8), which suggests that spatially varying selection resulting from environmental heterogeneity strongly favours the coexistence of alternative life-history strategies and thus the underlying genetic polymorphism. As expected, in additional models accounting for strong overdominance in survival (medium density conditions, Fig. S9) or negative frequency-dependant effect on male success (Fig. S10), most combinations of parameter values led to a protected polymorphism.

Estimating total fitness, defined as the combination of egg-to-adult survival and reproductive success for each genotype and sex, showed that antagonistic pleiotropy translates into two mechanisms of balancing selection: overdominance and sexual antagonism (Fig. 5A). Overdominance naturally emerges from antagonistic pleiotropy, even in the absence of dominance for any given trait, which is particularly true when selection is strong (Fig. 5D). In the case of *C. frigida*, heterozygotes benefit from the combination of a survival advantage associated with *β* and increased reproductive success associated with *α*. Variation in the strength and direction of dominance between fitness components, as observed in our experiment, with for instance overdominance in *αβ* male survival and dominance of *β* allele in female larval survival (Fig. 2), considerably expanded the parameter space leading to overdominance and polymorphism maintenance (Fig. 5D). Sexual antagonism emerges from the sex-specific effects of each allele on survival and reproduction, even in the absence of sexual antagonism for each fitness component (Fig. 5E), and also expands the range of conditions allowing polymorphism.

## Discussion

Results from the experimental evolution trial coupled with simulations show that the inversion dynamics in *Coelopa frigida* are largely governed by an antagonistic relationship between viability and fecundity. Using individual-based simulations, we highlight that this antagonism is probably stronger in natural conditions and contributes to the widespread maintenance of inversion polymorphism in the wild. The antagonistic pleiotropy between growth and reproduction observed in *C. frigida* at the inversion corroborates the increasing evidence for trade-offs mediated by body size (48–50). Such trade-offs frequently emerge because a bigger size either requires prolonged development time or a faster growth rate, both mechanisms associated with a lower likelihood to reach the reproductive stage. For instance, artificial selection for large body size in the yellow dung fly (*Scathophaga stercoraria*) increases juvenile mortality, particularly in stressful environments, as well as development time, leading to mortality before reproduction due to winter frost (51). For a wide range of species, environmental factors impose such limits to development time. For example, in snow voles (*Chionomys nivalis*) the benefits of higher body size are counteracted by the need to reach adult size before the first snowfall (52, 53). Similarly, in many annual plants, delayed flowering allows investment in vegetative growth and a subsequently higher number of seeds, but is constrained by the duration of the reproductive window (24, 28, 54). In *C. frigida,* this effect is relevant given the instability of the wrackbed habitat due to tide and wind induced waves (here modelled as limited habitat availability). By disproportionately increasing mortality of the genotypes with larger reproductive success and longer development time, the effect of the environment is expected to exacerbate the trade-offs reported during the experimental trials.

When life-history trade-offs are affected by external factors, environmental heterogeneity in space and time strongly contributes to the maintenance of variation by locally favouring different life-history strategies. For instance, in the yellow monkeyflower *Mimulus* spp., spatial heterogeneity in wetness and seasonal variation alternatively favours alleles determining early-flowering/low fecundity or late-flowering/high fecundity (24, 28, 29). In *C. frigida,* wrackbed stability varies between locations and across seasons depending on the exposure of the beach to tidal and storm-induced waves. Such heterogeneity is expected to generate fluctuating selection regimes between a slow-development/high fertility strategy (*αα*) or a fast-development/low fertility strategy (*ββ*), leading to variations in inversion frequency, and enhancing the maintenance of polymorphism. This prediction is supported by field observations reporting temporal variation in genotypic proportions, with the *α* allele increasing in frequency in summer when the wrackbed was less frequently disturbed by storms (38). Geographic variation in genotypic proportions are also observed in natural populations of *C. frigida* and have been associated with environmental variations, such as air temperature, depth and temperature of the wrackbed and substrate composition (30, 31). Although those factors may correlate with the duration of the wrackbed stability, they are also known to modulate the genotype-phenotype relationships and therefore the associated fitness. Controlled experiments in *C. frigida* show that substrate composition, temperature and density affect the relationship between genotype and survival, development time, body weight and body size (GxE effect) (37, 41, 55, 56). In nature, this translates into geographic variation in adult size differences among genotypes and between the sexes (30, 40), which may modify the reproductive advantage associated with the *α* allele. In other words, local environmental conditions are expected to also affect the slope of the trade-off between survival and fertility in *C. frigida*. Many life-history trade-offs are probably modulated by environmental effects: for instance, genetic correlations have been shown to vary in direction and strength between environments (57). Spatio-temporal environmental heterogeneity has thus a bivalent effect, by causing not only fluctuations in selection but also in the intensity of the trade-off. Both of these aspects interact and favour the maintenance of genetic variation.

In addition to environmental heterogeneity, polymorphism generated by antagonistic pleiotropy is predicted to be enhanced by different trade-off intensities for each sex (23). Sex-specific selection on different fitness components frequently occurs in nature since optimal age/size at maturity or the physiological state to achieve high fertility often differs between sexes (14, 58, 59). *Coelopa frigida* provides an empirical case of antagonistic pleiotropy whereby the strength of selection (but not the direction) differs between sexes for each fitness component. Differences between genotypes in size, development time, fertility and survival are stronger for males than for females. Within a range of realistic parameters, sex-specific effects, even without antagonism for each fitness components, results in sexual antagonism for total fitness, therefore strongly protecting polymorphism (23). Moreover, even without sexual antagonism in total fitness, our simulations suggest that sex-specific selection varying between components of fitness delays the fixation of alleles (Fig S11). While this mechanism cannot on its own protect long-term polymorphism, the slower rate of allelic fixation might allow additional factors, such as environmental heterogeneity, to prevent allele fixation (26, 60).

Overdominance emerges as a by-product of antagonistic pleiotropy, since the total fitness of heterozygotes is generally higher than both the fitness of homozygotes: *αβ* heterozygotes reach the highest fitness by combining a reproductive advantage due to large size (larger than *ββ*) and a survival advantage in natural conditions due to a shorter development time (shorter than *αα*). The parameter space leading to emergent overdominance is even more extended when dominance varies between fitness components, especially in the case of dominance reversal, that is when advantageous alleles for each component of fitness are dominant (22, 27). Since the reversal of dominance between fitness components has only been explored theoretically so far, it is still unknown how common it is in nature and what genetic architecture may underlie it. Our data still suggests that, in *C. frigida,* dominance at the inversion locus does vary in strength and direction between survival and fertility, as well as between sexes. These findings are in line with recent studies in other species showing that dominance can vary depending on the sex (14, 15, 61), and that multivariate traits, such as total fitness or complex traits, often involve the combination of trait-specific dominance (62–64). Overdominance may also emerge from the genetic architecture of the trait under antagonistic pleiotropy: inversions are frequently associated with deleterious or lethal effects, either because the breakpoints disrupt a gene, or because they contain recessive deleterious mutations which can only be purged by recombination and purifying selection when they occur at the homozygous state (33, 65–67). The generally higher egg-to-adult viability of heterozygotes, which is enhanced in stressful (high-density) conditions, suggests that the two haplotypic rearrangements of *C. frigida* may contain different clusters of deleterious mutations, a result supported by intra-and inter-population crosses (36). Such a linked genetic load may explain why the rarest *αα* genotype exhibits the highest deficit compared to Hardy-Weinberg expectations in nature (30) and why the *αα* genotype has the lowest viability in laboratory conditions. Since *α* is less frequently found at the homozygous state than the *β* allele in nature, the purging of deleterious mutations linked to the *α* allele might be more limited than for the *β* allele. Overdominance may hence be linked to two mechanisms, antagonistic pleiotropy and associated genetic load, which could enhance each other. Overdominance emerging from antagonistic pleiotropy may generate an excess of heterozygotes, which in turn limit the purging of the different recessive deleterious mutations associated with the two haplotypes. Such a sheltering of the genetic load associated with the inversions may in turn reinforce overdominance.

Inversions are frequently reported as polymorphic and maintained over long evolutionary timescales by balancing selection (34, 35, 68, 69). While one of the reasons could be the genetic load linked to the lack of recombination, we argue that antagonistic pleiotropy may also be a more important feature of the inversion systems than previously acknowledged. In fact, the particular architecture of inversions leads them to behave as a single-locus because of the stark reduction of recombination between rearrangements, but they are composed of dozens to hundreds of genes, sometimes interacting in a so-called “supergene” complex (33, 70, 71), where combinations of alleles at different genes lead to highly differentiated phenotypes. Inversion haplotypic rearrangements can thus have large, pleiotropic effects on complex phenotypes, which will be under various selective pressures, possibly in opposing direction (34). For example, in the longwing butterfly *Heliconius numata* an inversion polymorphism determining wing colour pattern is under opposing pleiotropic selection, with survival under positive frequency-dependent selection and reproduction under negative frequency-dependent selection (72). Likewise, in the ruff *Calidris pugnax* a polymorphic inversion that determines male reproductive morphs carries inverted alleles with lethal effect on survival but positive effects on testis size, indicating a possibly higher reproductive success, which is also under frequency-dependent selection (73). In the rainbow trout *Oncorhynchus mykiss,* alternate reproductive tactics (semelparous or iteroparous) are determined by a large polymorphic inversion, which involves a trade-off between reproductive output and longevity as well as a possible frequency-dependent effect (74). The accumulating evidence from the literature combined with our results on *C. frigida* thus indicate that contrasted effects of inversions on different fitness components may represent a non-negligible process involved in the maintenance of inversion polymorphism. We thus have to consider antagonistic pleiotropy and the effect of a wide spectrum of fitness traits on pleiotropic interactions more carefully when studying the evolutionary importance of structural genomic variants, that are currently increasingly documented to underlie complex phenotypes (34, 35, 68).

## Conclusion

Based on experimental evolution and extensive simulations, we demonstrate that a large chromosomal inversion is associated with a trade-off between larval survival and adult reproduction in the seaweed fly *C. frigida*. Such antagonistic pleiotropy contributes to the maintenance of this inversion polymorphism in nature. Earlier theory had considered antagonistic pleiotropy to play only a minor role in balancing selection because, taken on its own, it maintains polymorphism only under very restricted conditions. Yet, here, we provide a strong empirical case showing that this process occurs in nature, and more importantly, how antagonistic pleiotropy may realistically interact with other sources of balancing selection to further enhance the likelihood of persistent genetic polymorphism. For instance, the intensity of the trade-off varies with the sex and with the environment, which leads, respectively, to sexual antagonism and spatial/temporal fluctuations in allele frequencies, two mechanisms favouring the persistence of genetic variation. Similarly, overdominance naturally emerges from antagonistic pleiotropy, particularly when dominance varies between traits, as observed in *C. frigida.* Such emerging overdominance may lead to heterozygote excess, which in the case of an inversion could favour genetic load in homozygotes and enhance overdominance over time. Overall, our findings will stimulate research to reconsider antagonistic pleiotropy as a key part of multi-headed balancing selection processes that maintain genetic variation in nature.

## Materials and Methods

### Experimental evolution *in vivo*

Starting with two populations of wild *Coelopa frigida*, collected along the coast of the Gulf of St. Lawrence in Québec (Canada), we set up 16 replicates of experimental evolution (2 populations * 2 substrates * 4 replicates). The experiment is described in details in the supplementary methods. Briefly, each replicate was kept isolated and maintained for 5 generations under semi-natural conditions, controlling for temperature, density and substrate. Half of the replicates were kept on Laminariaceae and the other half on Fucaceae, the two main substrates on which *C. frigida* flies are naturally found in this region (Fig. 1B). To disentangle the fitness components, the generations were non-overlapping and we explicitly separated the growing phase from the reproductive phase. For reproduction, the pool of adults was left overnight at 25°C on a layer of seaweeds, either Laminariaceae or Fucaceae. After 16-20h, all eggs were collected in salt water and adults were preserved in ethanol. A subset of 1000 eggs was transferred to a controlled mass of seaweeds to start the growing phase of the next generation, and a subset of eggs was preserved in ethanol or RNAlater for subsequent genotyping. Upon emergence, adults were collected every day by aspiration and kept in conditions favouring high survival but without reproduction (5°C, dark, with a solution of Mannitol 0.5%). The frequency of the inversion and the proportions of the three inversion genotypes (*αα, αβ, ββ*) were estimated by genotyping adults (40-95 per replicate) and eggs (28 - 51 per replicate), using a diagnostic SNP assay (30). The inversion frequency was analysed with generalized linear mixed model for binomial data taking into account the identity of the replicate as random factor.

### Estimates of fitness

Relative survival rate of each genotype (and sex) was calculated by comparing the genotypic proportions in adults to those same proportions observed in the eggs. The ratio of these values is expected to be 1 if all genotypes survive equally. Similarly, we calculated the ratio of genotypic proportions in the eggs according to expected Hardy-Weinberg proportions based on the previous generation. This ratio is expected to be around 1 if mating occurs randomly and if all genotypes reproduce equally. Differences between genotypes in survival rate or in the deviation from random mating in the eggs were tested with a linear mixed model and post-hoc *t*-test. Development time was measured at generation 5 on 757 adults and differences between genotypes and sex were tested by comparing generalized linear mixed models based on a Poisson distribution. Female fecundity was assessed using 90 females by counting the number of eggs laid by each female within a 12 – 24 h period and then tested with linear models. All statistical analyses were performed using the software R 3.5.0 (75). Additional details are provided in the supplementary methods.

### Simulating evolution *in silico*

To evaluate how fitness differences between genotypes based on a different investment in the trade-off between survival and reproduction modulates the evolution of the inversion frequency, we developed an individual-based model inspired by the results of the experimental evolution trials and by the biological characteristics of *C. frigida* (Fig. 3A). The model is described in detail in the supplementary methods following the ODD protocol (76). Briefly, generations were non-overlapping and the time step was one generation. Each egg goes through a growing period, during which its survival is determined probabilistically by its sex and genotype, and then matures into an adult if its development time is shorter than the duration of its habitat. To model heterogeneity in the duration of habitat availability, this value was randomly-drawn for each individual in a uniform distribution determined by two parameters: mean duration and variability. Adults then go through a reproductive event. Each male can mate several times with a probability influenced by its genotype and based on the genotype proportions of all mature males. All females reproduce once and lay a number of eggs depending on their genotype. Egg genotype is determined by Mendelian inheritance from parental genotypes. A subset of *K* eggs initiates the next generation, mirroring the census made in the experiment or a limited carrying capacity in the wild. At each generation, we recorded the proportions of the three genotypes in the eggs and the adults. The model was implemented in Rust 1.32.0 (9fda7c223 2019-01-16) and the code and parameter files are available online (https://github.com/enormandeau/coelopa_fastsim).

The model outcomes were then analysed at three levels. (i) First, we ran the model for 5 generations with parameters drawn from the experiment while varying the relative male reproductive success. The fit of each simulation to empirical data was quantified by computing the normalized root-mean-squared error (nRMSE) for each genotypic proportion from generation 1 to 5. (ii) To simulate evolution in a natural population, we then ran the model with larger K (10 000) for 200 generations and included variation in environmental parameters. The equilibrium in genotypic proportions was compared to the frequencies observed in nature and we explored the combination of parameters possibly influencing the frequency of the inversion. We tested the role of density by exploring several sets of survival values based on previous laboratory studies (37) (Table S3). For relative male reproductive success, the full range of parameters was explored because this parameter could not be estimated empirically and possibly varies between populations with natural variation in adult size (30, 55). Habitat availability was set between 7-20 days, *i.e.* the range of development time found in our experimental conditions, to estimate the interplay between those two factors. Yet, in natural conditions, both the range of wrackbed availability and development time are expected to be wider. Development time varies with density and temperature, although the ordering of emergence between the three male genotypes remains unchanged. Wrackbeds are expected to be removed cyclically by spring tides, so the actual availability would be slightly less than 14 or 28 days, but they are sometimes observed to last shorter because of storms (47). (iii) Finally, the backbone of the model was used to theoretically explore the range of conditions under which antagonistic pleiotropy could maintain polymorphism at evolutionary time-scales when combined with sex-specific effects and dominance, (*K*=10 000, 500 generations). Total fitness was calculated as the product of the relative survival rate and reproductive success, for each sex and genotype.

## Acknowledgments

We are very grateful to C. Babin, B. Labbé and L. Logacheva for help in the field and with raising and genotyping the flies. We would like to thank F. Larochelle, L. Johnson and J-C Therrien for their support in establishing the fly rearing conditions. We would also like to thank E. Berdan for valuable discussion about *Coelopa frigida* biology and the inversion dynamics. M. Laporte and H. Cayuela provided constructive comments on the manuscript. This research was supported by a Discovery research grant from the Natural Sciences and Engineering Research Council of Canada (NSERC) to L.B., by the Canadian Research Chair in genomics and conservation of aquatic resources held by L.B. and by the Swedish Research Council grant 2012-3996 to MW. C.M. was supported by a post-doctoral fellowship from the FRQNT and FRQS.

## Data deposition

Simulation code is available at https://github.com/enormandeau/coelopa_fastsim with parameters files; Experimental data will be deposited on Dryad.

## Authors’ contributions

C.M., M.W. and L.B. designed the research. C.M. performed the research and analyzed the data. E.N., C. M and V. L. developed the model. C.M., V. L., E. N., M.W. and L.B. wrote the manuscript.

