## Supplementary Materials for "Balancing selection via life-history trade-offs maintains an inversion polymorphism in a seaweed fly"

### *Appendix of methods*

#### Experimental evolution

- Field sampling

A large number of *Coelopa frigida* adult flies were sampled in their natural habitat of decomposing seaweed (wrackbed) at two locations along the Canadian Atlantic coast, in May 2016 at Cap Espoir, Québec (CE: 48.43087, -64.32778) and in June 2016 at Kamourasksa, Québec (KA: 47.56294, -69.87375). In Cap Espoir, the wrackbed was dominated by Laminariaceae, while at Kamouraska the wrackbed was mostly composed of Fucaceae (1). The wild-caught adult flies constitute the generation 0 of the experimental evolution procedure. They were kept for a few days at 5°C in the dark, in plastic bags with a cotton soaked in 0.05% Mannitol; conditions which are known to allow high survival but no reproduction (2). The two populations (KA & CE) were kept separated and represented two distinct experimental lines. Natural seaweed from the families Laminariaceae and Fucaceae were collected (either attached to the substrate or floating in the sea) in Métis sur Mer, Québec (48.66408, -68.07221) first in May 2016 and then repeatedly over the course of the experiment. Seaweed were transported to Laval University and frozen for subsequent use as substrate in the experiment. Freezing allows a good conservation of the seaweed and destroys any eggs, parasites or larvae.

- Wild seaweed fly collection and initiation of the experimental evolution trial

Wild adult flies (KA: 303 females, 218 males, CE: 396 females, 570 males) were left overnight (for 16 hours) at 25°C in 5 boxes per population with a substrate of natural seaweed including both Fucaceae and Laminariaceae, conditions known to favour reproduction and egg laying behaviour. After 16 hours, all adults were captured and preserved in ethanol for subsequent genotyping. Eggs were collected by flooding the seaweed substrate with 3% salt water (equivalent to sea water) in each box, and pooled per population. A subset of the eggs was preserved in ethanol and RNAlater for subsequent analysis. Egg density in the salt water solution was counted over a grid in three subsets of 3 mL each. A volume corresponding to an estimated 1000 eggs was filtered over a dark piece of linen, constituting generation 1. This was repeated 10 times to start 10 replicates per population at generation 1.

- The main experimental evolution trial

At each generation, the linen with 1000 eggs from a given replicate was deposited in a raising box (one box per replicate) on top of the seaweed substrate and kept until all adults emerged.

For all replicates, raising boxes were identical plastic boxes measuring 12x8x18 cm. The boxes were closed with a hermetic lid perforated with four holes filled with sponge to allow air exchange but preventing adult flies to escape. Stones and sand were put in the bottom of the box to facilitate drainage. The 1000 eggs started on 500g substrate with subsequent addition of 200g of seaweeds after approximately 6 days. This procedure ensured a controlled larval density between replicates. To make the substrate, seaweeds were thawed at 5° for 24h-48h and cut into large pieces for Laminariaceae or grinded into small pieces for Fucaceae. This difference in the scale of grinding reflects the inherent differences between the two kinds of seaweed and their usual state in natural wrackbeds. Half of the replicates constituted of 90% of Laminariaceae and 10% of Fucaceae, and the other half with 90% of Fucaceae and 10% of Laminariaceae. All replicates were kept in chambers controlled for temperature (25°C), humidity (50%) and light duration (12h).

After approximately 7-8 days, the first adults started to emerge. From that day, adults were collected daily by aspirating all the flies with a small insect vacuum. Flies were kept separate for each replicate at 5°C in the dark in plastic bags closed hermetically and supplemented with cotton soaked in 0.05% Mannitol. When all adults emerged, we initiated the following generation with the same procedure as described for generation

0, except that each replicate was kept separate. In other words, free mating was allowed between all adults emerging from one replicate and 1000 eggs were gathered for the next generation. The remaining eggs and reproductive adults were preserved for subsequent genetic analysis.

Only 16 replicates (8 replicates per population) were kept until generation 5 due to stochastic crashes in population size experienced by some. Furthermore, to avoid founder effects, we allowed reproduction and the initiation of a subsequent generation only for those replicates in which more than 100 adults emerged per generation.

- Monitoring of the inversion frequencies over time

Genomic DNA was extracted individually with a lysis procedure for 96 adults per population (48 females/48 males) at generation 0, 48 adults per replicate (24 females/24 males) at generations 1 and 5 for all replicates, and 48 adults per replicate (24 females/24 males) at generation 2, 3 and 4 for four replicates. Genomic DNA was also extracted individually for 48-192 eggs per replicate at all generations for the same four replicates, with a lysis procedure in a smaller volume (20µL). DNA from each egg or from each adult was then genotyped for the inversion with previously described diagnostic SNP markers with PCR amplification and digestion by restrictions enzyme (1). While DNA extractions from adults produced good amounts of quality DNA (frequency estimates are based on over 40 adults per replicate/per generation), as it was more challenging with small eggs. Thus, we extracted as many eggs per replicate as possible aiming for at least 30 eggs genotyped per replicate per generation.

- Statistical analyses

Variation of genotype proportions between generations 1 and 5 were analysed with a generalized mixed model with a logistic link function for binomial data in the R package *lme4* (3) and *lmerTest* (4). The response variable was the number of individuals carrying *vs.* not carrying the given genotype, the random effect was the identity of the box replicate, and the fixed explanatory variables were generation and substrate or population.

Egg-to-adult survival rates were calculated for both sexes combined, as well as for each sex separately, by dividing the observed frequencies of the adults by the frequencies in the eggs. In fact, if all genotypes survived equally, frequencies in eggs and adults are expected to be the same and, for all genotypes, the ratio of frequencies should be centred around 1. Deviation from this expectation thus represents an increase or a decrease in egg-to adult survival relative to the other genotypes. Relative survival rates could be estimated in 32 boxes (i.e. all replicates and generations for which we obtained frequencies for the adults and in the eggs).

We compared frequencies observed in the eggs and the frequencies expected under random mating, i.e. Hardy-Weinberg proportions of the previous generation with a $X^2$-test in 18 boxes (i.e. all replicates and generations for which we obtained frequencies of the eggs and adults of the previous generation), and then with a meta-analyses on this set of p-values using weighted Z-method in the R package *metap* (5). If mating occurs randomly and if all genotypes have equal fecundity, the ratio of observed frequencies in the eggs over expected frequencies should be 1. Deviations from this expectation thus represent an estimate of the relative reproductive success between genotypes.

Variation in relative survival and relative reproductive success was tested with a linear mixed model, with genotype and sex as fixed effects, and box identity as a random effect with the R packages *lme4* (3) and *lmerTest* (4); followed by a paired post hoc t-test (corrected following (6)). We also explored generation, population and substrate as fixed effects, although not all interactions could be tested given the small sample size.

#### Follow-up measurements of life-history traits

- Development time

Once the 5th generation of experimental evolution was reached, we recorded individual development time from egg to adult. Variation in development time was tested with a generalized linear mixed model based on a Poisson distribution, with genotype, sex or substrate as fixed effects and box identity as a random effect, with individual flies being the unit of replication. Each factor and interaction was added sequentially and nested models were built with the R package lme4 (3) and compared to each other with a $\chi^2$-test to determine whether the additional factor significantly improved the explicative power of the model.

- Female fecundity

We carried out a follow-up experiment to test whether female fecundity, measured as the number of eggs in the first clutch, differed between genotypes. We established a laboratory population with the same method as before, using eggs laid by wild-caught females from Kamouraska, Québec, which we raised under the same conditions as described for the previous experiment (25°C, substrate of 90% Laminariaceae- 10% Fucaceae, low density). After 7 days, pupae were collected every day and kept isolated in a 2mL tube with cotton soaked in 0.05% Mannitol. Upon emergence, adults were kept for a few days at 5°C in the dark. A total of 161 pairs of one male and one female were formed. On the day of the experiment, each pair was transferred into a small box (approximately 5x5x5 cm) with a mix of Laminariaceae and Fucaceae, and left overnight (for 16h) at 25°C (identical conditions as used in the previous experiment for mating and egg laying). After 16h, if a clutch of eggs was observed, it was collected and preserved on a dark-linen in a 1.5mL tube of RNAlater, and the parents were preserved in ethanol. If no eggs were observed, the pair was kept in the same experimental box and the eggs were checked again after an extra 8h. We did this to maximise the number of females for which we could count eggs (eggs after 16h: 69 females, eggs after 24h: 120 females). We genotyped 90 females for the inversion using the method described above and counted their eggs under a binocular magnifier (Zeiss Stemi 2000C). Variation in the number of eggs per female in relation to female genotype (αα, αβ, ββ) and time of laying (16h/24h) was analysed with a linear model and post hoc pairwise t-test (adjusted following(6)). The fact that eggs were laid after 16h or 24h did not significantly affect the number of eggs per female ($F_{1,88}$=0.09, p=0.76), or the interaction with genotype ($F_{2,84}$=1.96, p=0.15), and we therefore pooled the data for all subsequent analyses to derive female fitness parameters in the model.

#### Simulation model

- Overview (Fig. 3A, Table 1, Table S7)

*Purpose:* The purpose of the model was to evaluate how differences between genotypes in reproductive success and survival affect the evolution of the inversion genotype frequency in the experiment or in the wild and, more generally, the maintenance of polymorphism. *State variables and scales:* The model is based on individuals, which belong to the same population. Individuals are characterized by three state variables: stage (egg or adult), sex (male or female), inversion genotype (αα, αβ, ββ) and three traits: egg-to-adult survival, development time and reproductive success, which are dependent on sex and inversion genotype. Inversion alleles are inherited in a Mendelian fashion. *Process overview and scheduling:* The model consists of non-overlapping generations. Each time step is one generation. At each time step, two phases are processed in this order: reproduction of generation n-1, resulting in the population of eggs of the generation n, and egg-to-adult growth, resulting in the population of adults of generation n.

- Design concepts

*Basic principles:* The basic principle by which the model is constructed is a trade-off between two components of fitness, egg-to-adult survival and reproduction, which vary with the inversion genotype. Some scenarios include complementary features such as sex-specific parameters, trait-specific dominance including overdominance, frequency-dependence and environmental variation. *Adaptation:* Individuals have three adaptive traits, which are fully determined by their sex and genotype: egg-to-adult survival, development time and reproductive success. *Emergence:* Genotype frequencies at each generation, and thus inversion allelic frequencies, emerge as properties of the population from the relative fitness of the individuals. *Stochasticity:* All individual traits (survival, development time, relative reproductive success) and environmental variation (duration of habitat availability) are interpreted as probabilities, or are drawn from empirical probability distributions. Demographic stochasticity, somehow equivalent to a limited charge capacity, was included by randomly picking a subset of K eggs at the beginning of the growth phase. *Interaction:* Reproduction is modelled explicitly in relation to the number of available males and genotype-specific reproductive success. Duration of habitat availability interacts with individual development time and affects egg-to-adult survival. *Observation:* Genotype frequencies in the eggs and in the adults are the variable that we monitored across time steps.

- Details

*Initialization:* The model is initialized with $N_0$ adults at a sex-ratio of 50:50 and genotype proportions as observed in wild populations, i.e. in the experimental generation 0. $N_0$ is the product of K, the number of eggs kept for growth and $S_0$, the absolute survival rate.

*Submodels - Reproduction:* At each reproductive step, a pool of reproductive males was generated based on the distribution of adult males $\{N_{\alpha\alpha\text{-}m} ; N_{\alpha\beta\text{-}m} ; N_{\beta\beta\text{-}m}\}$, corrected by genotype-specific male reproductive success following $\{ N_{\alpha\alpha\text{-}m} * T_{\alpha\alpha\text{-}m}; N_{\alpha\beta\text{-}m} * T_{\alpha\beta\text{-}m}; N_{\beta\beta\text{-}m} * T_{\beta\beta\text{-}m} \}$. This allowed modelling male reproductive success as a relative availability and propension for mating and took into account that males could reproduce many times in this species. For females, mixed paternity has been reported for *C. frigida* in only 5-10 % of the females (7), thus for simplicity, all females reproduced only once in the model. All females were considered successively in a random order. For each female, a male partner was randomly drawn from the pool of reproductive males, and the number of eggs laid by the pair was determined by the product between the number of eggs by a female and genotype-specific female fertility. Each egg inherited three state variables: (i) sex, which is determined by chance with no bias (sex-ratio= 0.5), (ii) genotype, which is assigned by randomly drawing one allele from the mother, and one allele from the father, (iii) development time, which was calculated as the cubic root of three values randomly-drawn from a uniform distribution whose mean and range was determined by sex and genotype. This distribution was chosen as the one fitting best the measured experimental data of the development time. At the end of the reproductive step, a subset of randomly-picked K eggs proceeded to the growth step. This feature mimicked the experimental procedure and ensured a constant population size, being somehow equivalent to a carrying capacity.

*Submodels – Growth:* The outcome (survival or death) of each egg entering the growth phase was determined by chance with a survival probability determined by the product between global egg-to adult survival rate and relative genotype-sex specific survival. This step represents a form of "intrinsic mortality", observed even in favourable laboratory conditions. Yet, in the wild, *C. frigida*'s habitat is known to be temporary: the wrackbed can be removed monthly or bi-monthly by tides, and occasionally by storms (8). To take this environmental effect into account, each surviving individual was attributed a habitat availability duration, randomly-drawn from a uniform distribution centred on $A_{mean}$, with width $A_{var}$. Variation in the duration of habitat availability between individuals can be interpreted as variation in the moment at which the egg was laid, as well as heterogeneity between wrackbeds. If development time exceeded the duration of habitat availability, the individual did not proceed to the adult stage. For models that do not consider such environmental effect, $A_{mean}$ was taken as a very large value (30 days) and thus did not affect the total egg-to-adult survival.

- Simulated scenario and analysis

*A mirror of the experimental evolution:* To understand the dynamics of genotypic frequencies in the experimental system and to infer the parameters that could not be measured experimentally, such as male mating success, we ran the model over 5 generations, with K=1000 eggs, all parameters set to the measured experimental values and 30 simulations per set of parameters. We then explored several scenarios, (1) setting αα male mating success at 1 and varying male mating success of ββ between 0.05-1, αβ mating success being the mean of αα and ββ values (co-dominance), (2) setting αα and αβ mating success at 1 and varying male mating success of ββ between 0.05-1 (dominance), (3) setting αα male mating success at 1, and determining male mating success of αβ and ββ males as a function of αα-males frequency ($F_{\alpha\alpha\text{-m}}$) and two parameters (*FDC*, a coefficient of frequency dependence, and $t'_{\beta\beta\text{-}m}$, a fixed parameter) with the following equations:

$$T_{\beta\beta-m} = t'_{\beta\beta-m} \cdot \left(1 - FDC \cdot (1 - \mathrm{F}_{\alpha\alpha-\mathrm{m}})\right)$$

$$T_{\alpha\beta-m} = \frac{1 + t'_{\beta\beta-m}}{2} \cdot \frac{\left(1 - FDC \cdot (1 - \mathrm{F}_{\alpha\alpha-\mathrm{m}})\right)}{2}$$

For all scenarios, we compared the evolution of genotype proportions in the eggs and the adults over 5 generation to the empirically-observed evolution scenario. The fit of each simulation to empirical data was quantified by computing the normalized root-mean-squared error (nRMSE) for each genotypic proportion for generations 1 to 5. The average nRMSE over the 6 variables (αα/αβ/ββ proportions in the eggs and the adults), and over the 30 replicates, was taken as an index of fit, with the best predicting scenarios having the smallest values. Difference of mean nRMSE between the best scenarios was tested with a t-test based on the 30 replicates, corrected following (6). Visualization of the evolution of frequencies was made with the R package *ggplot2* (9)

*A model of natural populations:* To expand the scope of the model to natural populations, the simulations were run over 200 generations with K= 10000 and 30 replicates per set of parameters. The null model was based on the parameters estimated in the experiment (and for male reproductive success, on parameters inferred from the best model fitted over 5 generations as described above). We then explored different sets of parameters that could represent a better approximation of natural conditions than experimental values (intermediate male reproductive success, limited substrate availability) or that present natural environmental variability (density, duration of habitat availability). This was done by following three scenarios that vary one parameter at a time and a scenario varying two parameters: (i) A scenario exploring survival variation with values estimated at medium and high density by Butlin *et al.* (10) (ii) A scenario varying male mating success of ββ between 0.05 and 1 in a co-dominance scenario, (iii) A scenario taking into account the length of habitat availability by varying $A_{mean}$ between 7 and 15 days, (iv) a scenario setting male mating success of ββ at an intermediate value (0.5) and varying the duration of habitat availability from 7 to 15 days with two survival conditions, low (experimental) density and medium density. Genotypic proportions of the adults at the 200$^{th}$ generation were compared to values observed in wild populations (1, 11) and at the 5$^{th}$ generation of our experiment. All simulations had reached stable genotypic proportions before the 100$^{th}$

generation. The proportion of the three genotypes under each scenario was visualized in ternary plots built with the R package *ggtern* (12). Next, for the parameters that are more likely to vary in natural populations (male relative success, duration of habitat availability, density, variability in the duration of habitat availability), we explored under which combinations of realistic parameters polymorphism was maintained after 200 generations, what was the mean frequency of the inversion at equilibrium after 100 replicates, which portion of the parameter space lead to polymorphism and whether overdominance or sexual antagonism emerged for total fitness.
Total fitness ($W_{XX-s}$) was calculated as the product of relative survival rate and reproductive success, for each genotype (*XX, being αα, αβ or ββ*) and sex (*s, being f or m*) and for each allele, as:

$$W_{xx-s} = S_{xx-s} \cdot T_{xx-s}$$

$$w_{\alpha-s} = w_{\alpha a-s} + \frac{1}{2} \cdot w_{\alpha\beta-s}$$
$$w_{\beta-s} = w_{\beta\beta-s} + \frac{1}{2} \cdot w_{\alpha\beta-s}$$

Overdominance for total fitness corresponded to cases in which:

$$w_{a\beta-s} > w_{\alpha\alpha-s} \;\&\; w_{a\beta-s} > w_{\beta\beta-s}$$

Sexual antagonism for total fitness emerged in cases under which:

$$w_{a-m} > w_{\beta-m} \;\&\; w_{a-f} < w_{\beta-f} \qquad \text{or} \qquad w_{a-m} < w_{\beta-m} \;\&\; w_{a-f} > w_{\beta-f}$$

*Generalization:* To test more generally how polymorphism can be maintained by antagonistic pleiotropy in interaction with dominance/overdominance and sex-specific effects, we ran the same model based on a trade-off between survival and reproduction over 500 generations with K=10000 and 100 replicates. These simulations explored the whole theoretical parameter range for survival and reproduction (Table S7), with either various scenarios of dominance, coded by the parameters *Hs*/*Ht*, or sex-specific effects with independent values for *Sm/Tm* and *Sf/Tf* ranging between 0 and 1. We surveyed the proportions of the simulations that led to maintenance of polymorphism vs. the simulations in which one of the genotypes got fixed, as well as the emerging mechanism at the level of total fitness (overdominance in one/both sex, sexual antagonism). Initial proportions were set to Hardy-Weinberg proportions, with the frequency of α being 0.5. The main features of the models remained the same, except that the effect of habitat availability/development time was not taken into account. In fact, the environmental effect on genotype-specific survival can be generalized in the survival parameter.

### *Supplementary tables and figures*

#### Evolution of genotypic proportions in the experimental evolution

**Table S1: Number of samples genotyped and proportions of the three genotypes in the experiment.**
Wild stands for wild-caught flies, *i. e.* the founders, generation 0. L stands for 90% Laminariaceae, 10% Fucaceae; F stands for 90% Fucaceae, 10% Laminariaceae.

| | | | | | Adults | | | | Eggs | | |
|---|---|---|---|---|---|---|---|---|---|---|---|
| **population** | **replicate** | **generation** | **substrate** | **N** | **Proportions** | | | **N** | **Proportions** | | |
| | | | | | **αα** | **αβ** | **ββ** | | **αα** | **αβ** | **ββ** |
| | wild | 0 | | 94 | 0.09 | 0.54 | 0.37 | | | | |
| | CE01 | 1 | L | 47 | 0.21 | 0.60 | 0.19 | 51 | 0.18 | 0.61 | 0.22 |
| | CE02 | 1 | L | 46 | 0.24 | 0.46 | 0.30 | 51 | 0.18 | 0.61 | 0.22 |
| | CE03 | 1 | L | 42 | 0.24 | 0.64 | 0.12 | 51 | 0.18 | 0.61 | 0.22 |
| | CE04 | 1 | L | 42 | 0.21 | 0.40 | 0.38 | 51 | 0.18 | 0.61 | 0.22 |
| | CE07 | 1 | F | 48 | 0.08 | 0.67 | 0.25 | 51 | 0.18 | 0.61 | 0.22 |
| | CE08 | 1 | F | 46 | 0.09 | 0.67 | 0.24 | 51 | 0.18 | 0.61 | 0.22 |
| | CE09 | 1 | F | 40 | 0.18 | 0.50 | 0.33 | 51 | 0.18 | 0.61 | 0.22 |
| | CE10 | 1 | F | 41 | 0.15 | 0.59 | 0.27 | 51 | 0.18 | 0.61 | 0.22 |
| | CE01 | 2 | L | 48 | 0.31 | 0.46 | 0.23 | 43 | 0.33 | 0.44 | 0.23 |
| | CE07 | 2 | F | 48 | 0.42 | 0.48 | 0.10 | 43 | 0.40 | 0.49 | 0.12 |
| CE | CE01 | 3 | L | 41 | 0.34 | 0.61 | 0.05 | 45 | 0.44 | 0.49 | 0.07 |
| | CE07 | 3 | F | 46 | 0.43 | 0.41 | 0.15 | 29 | 0.34 | 0.55 | 0.10 |
| | CE01 | 4 | L | 46 | 0.43 | 0.54 | 0.02 | 45 | 0.51 | 0.42 | 0.07 |
| | CE07 | 4 | F | 44 | 0.23 | 0.75 | 0.02 | 31 | 0.45 | 0.55 | 0.00 |
| | CE01 | 5 | L | 48 | 0.46 | 0.52 | 0.02 | 28 | 0.57 | 0.39 | 0.04 |
| | CE02 | 5 | L | 47 | 0.32 | 0.64 | 0.04 | | | | |
| | CE03 | 5 | L | 46 | 0.30 | 0.54 | 0.15 | | | | |
| | CE04 | 5 | L | 48 | 0.56 | 0.44 | 0.00 | | | | |
| | CE07 | 5 | F | 48 | 0.33 | 0.52 | 0.15 | 30 | 0.47 | 0.43 | 0.10 |
| | CE08 | 5 | F | 47 | 0.49 | 0.47 | 0.04 | | | | |
| | CE09 | 5 | F | 47 | 0.43 | 0.49 | 0.09 | | | | |
| | CE10 | 5 | F | 48 | 0.40 | 0.54 | 0.06 | | | | |
| | wild | 0 | | 95 | 0.05 | 0.44 | 0.51 | | | | |
| | KA21 | 1 | L | 48 | 0.06 | 0.52 | 0.42 | 43 | 0.16 | 0.58 | 0.26 |
| | KA23 | 1 | L | 47 | 0.06 | 0.62 | 0.32 | 43 | 0.16 | 0.58 | 0.26 |
| | KA24 | 1 | L | 40 | 0.13 | 0.50 | 0.38 | 43 | 0.16 | 0.58 | 0.26 |
| | KA25 | 1 | L | 43 | 0.07 | 0.63 | 0.30 | 43 | 0.16 | 0.58 | 0.26 |
| | KA26 | 1 | F | 48 | 0.13 | 0.56 | 0.31 | 43 | 0.16 | 0.58 | 0.26 |
| | KA27 | 1 | F | 48 | 0.04 | 0.48 | 0.48 | 43 | 0.16 | 0.58 | 0.26 |
| | KA28 | 1 | F | 47 | 0.21 | 0.51 | 0.28 | 43 | 0.16 | 0.58 | 0.26 |
| | KA30 | 1 | F | 42 | 0.07 | 0.48 | 0.45 | 43 | 0.16 | 0.58 | 0.26 |
| | KA21 | 2 | L | 48 | 0.17 | 0.73 | 0.10 | 60 | 0.22 | 0.58 | 0.20 |
| | KA27 | 2 | F | 46 | 0.20 | 0.65 | 0.15 | 36 | 0.22 | 0.53 | 0.25 |
| KA | KA21 | 3 | L | 42 | 0.43 | 0.48 | 0.10 | 34 | 0.44 | 0.44 | 0.12 |
| | KA27 | 3 | F | 45 | 0.33 | 0.64 | 0.02 | 30 | 0.40 | 0.47 | 0.13 |
| | KA21 | 4 | L | 44 | 0.43 | 0.50 | 0.07 | 31 | 0.42 | 0.52 | 0.06 |
| | KA27 | 4 | F | 42 | 0.33 | 0.64 | 0.02 | 35 | 0.43 | 0.49 | 0.09 |
| | KA21 | 5 | L | 47 | 0.43 | 0.51 | 0.06 | 33 | 0.36 | 0.58 | 0.06 |
| | KA23 | 5 | L | 44 | 0.57 | 0.36 | 0.07 | | | | |
| | KA24 | 5 | L | 41 | 0.34 | 0.61 | 0.05 | | | | |
| | KA25 | 5 | L | 48 | 0.25 | 0.73 | 0.02 | | | | |
| | KA26 | 5 | F | 43 | 0.47 | 0.47 | 0.07 | | | | |
| | KA27 | 5 | F | 47 | 0.45 | 0.49 | 0.06 | 45 | 0.40 | 0.47 | 0.13 |
| | KA28 | 5 | F | 43 | 0.42 | 0.53 | 0.05 | | | | |
| | KA30 | 5 | F | 42 | 0.43 | 0.55 | 0.02 | | | | |

**Table S2: Analysis of genotypic proportions**

**Generalized linear mixed model testing difference of genotypic proportions between generation 1 and 5 and either the effect of the population of origin or of the substrate.**

Values are the z-values of the GLMM and p-value in brackets. The interaction between substrate and population was not tested given the size of the sample size (n= 4 replicates per generation, per substrate and per population)

| | α | αα | αβ | ββ |
|---|---|---|---|---|
| generation | **10.6 (p<0.001)** | **8.8 (p<0.001)** | -1.4 (p=0.16) | **-7.9 (p<0.001)** |
| substrate | 1.1 (p=0.27) | 1.4 (p=0.16) | -0.49 (p=0.62) | -0.3 (p=0.78) |
| generation * substrate | -0.85 (p=0.40) | 1.5 (p=0.14) | 0.95 (p=0.34) | -0.55 (p=0.58) |

| | α | αα | αβ | ββ |
|---|---|---|---|---|
| generation | **8.2 (p<0.001)** | **6.9 (p<0.001)** | -1.3 (p=0.19) | **-6.5 (p<0.001)** |
| population | -3.6 (p<0.001) | -2.9 (p=0.004) | -0.9 (p=0.39) | 3.3 (p=0.001) |
| generation * population | 2.8 (p=0.005) | 2.6 (p=0.01) | 0.8 (p=0.41) | -2.3 (p=0.02) |

#### Egg-to adult relative survival in the experimental evolution

**Table S3: Estimates of relative egg-to-adult survival in the experiment and the litterature**

| | | $S_{\alpha\alpha\text{-}f}$ | $S_{\alpha\beta\text{-}f}$ | $S_{\beta\beta\text{-}f}$ | $S_{\alpha\alpha\text{-}m}$ | $S_{\alpha\beta\text{-}m}$ | $S_{\beta\beta\text{-}m}$ |
|---|---|---|---|---|---|---|---|
| Experimental data (32 replicates) | mean [sd] | 0.80 [0.36] | 1.01 [0.22] | 1.12 [0.7] | 0.88 [0.47] | 1.08 [0.29] | 0.96 [0.58] |
| | (normalized for comparison) | 0.79 | 1 | 1.11 | 0.81 | 1 | 0.88 |
| From Butlin *et al* (10) | Low density | 0.87 | 1 | 1.09 | 0.83 | 1 | 0.73 |
| | Medium density | 0.52 | 1 | 0.63 | 0.40 | 1 | 0.35 |
| | High density | 0.56 | 1 | 0.34 | 0.33 | 1 | 0.32 |
| From Gilburn *et al* (13) | mean [sd] | 0.64 [0.11] | 1.18 [0.11] | 0.91 [0.12] | 0.64 [0.11] | 1.18 [0.11] | 0.91 [0.12] |
| | (No sex-specific data) | 0.54 | 1 | 0.77 | 0.54 | 1 | 0.77 |

**Table S4: Analysis of relative survival rate**

**Linear mixed models testing the effect of genotype, sex, substrate or population on the**

Note: The effect of generation is driven by the variation in ββ survival (Fig. S2), which is somehow lower in the last generations. Yet, the effect of generation is difficult to test and may arise from stochasticity because sample size is small (4 replicates per generations) and because survival rates at generation 3 to 5 are not strictly reliable given the low absolute numbers of BB found in the eggs or the adults after generation 3.

| | **Df** | **F** | **P** |
|---|---|---|---|
| **genotype** | 2 | 4.7 | **0.01** |
| **sex** | 1 | 0 | 0.99 |
| **substrate** | 1 | 0.06 | 0.8 |
| **population** | 1 | 0.87 | 0.35 |
| **generation** | 1 | 0.7 | 0.4 |
| **genotype * sex** | 2 | 1.5 | 0.21 |
| **genotype * substrate** | 2 | 0.78 | 0.46 |
| **genotype * population** | 2 | 0.43 | 0.65 |
| **genotype * generation** | 2 | 11.2 | **<0.001** |

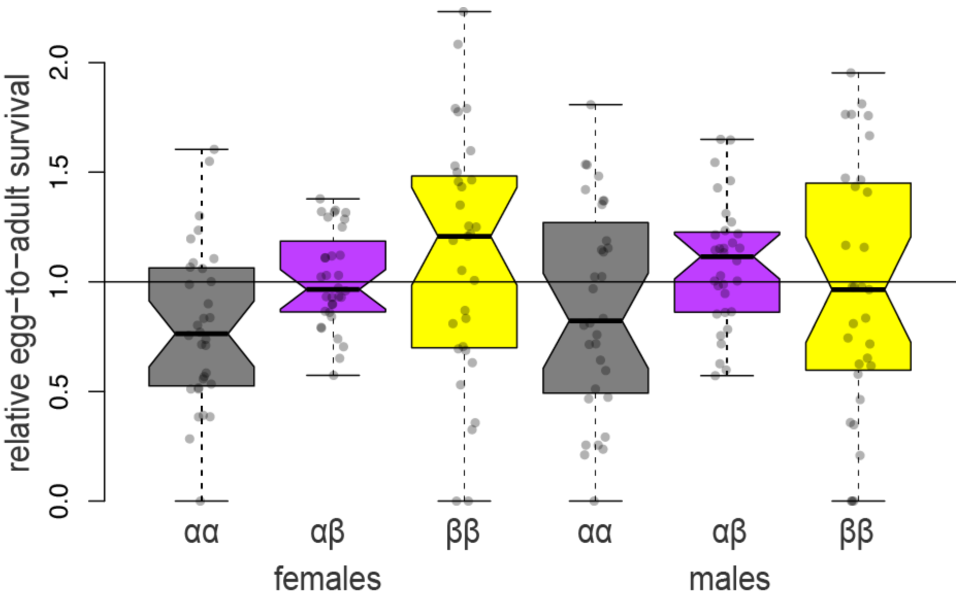

**Figure S1: Relative egg-to-adult survival rate per sex for each genotype**
Neither sex nor the interaction sex:genotype significantly affect the relative survival rate. Pairwise t-test do not show significant differences between mean survival rates because of the large variance heterogeneity.

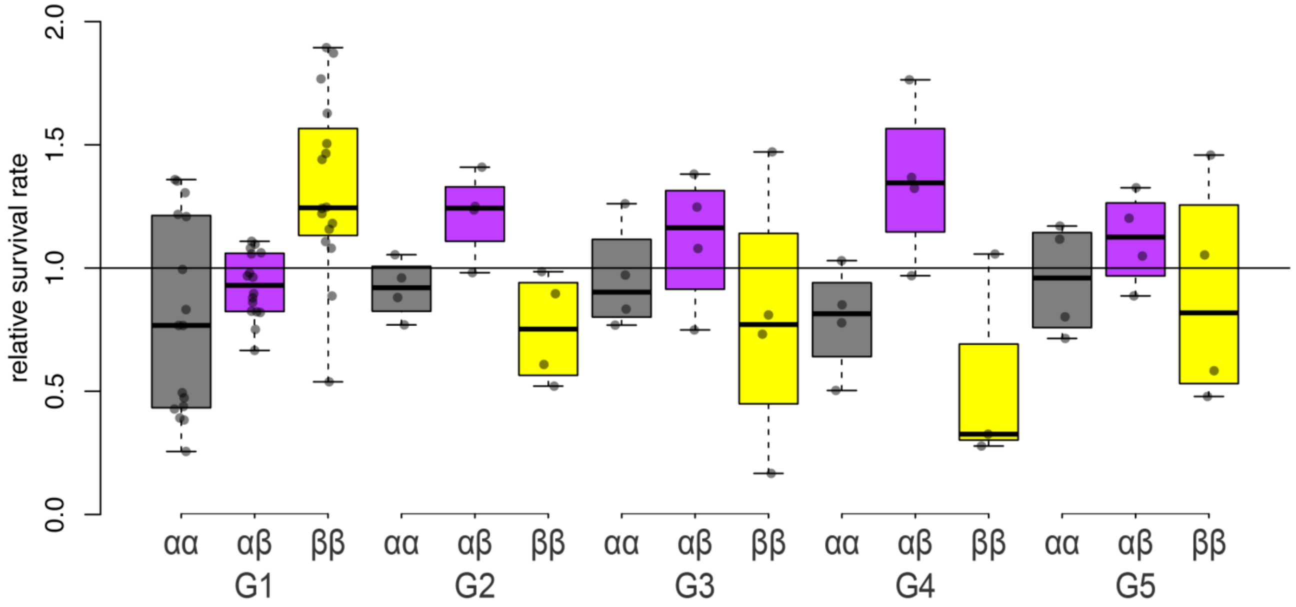

**Figure S2: Relative egg-to-adult survival rates per generation for each genotype**
Note: Although there seems to be variation in survival rate between generations and the test suggests a significant genotype*generation effect on survival rate (Table S2), there is no clear trend and much stochasticity may come from the fact that generations 2 to 5 include four replicates and survival rates at generation 3 to 5 are not extremely reliable given the low absolute numbers of BB found in the eggs or the adults after generation 4.

#### Development time at generation 5

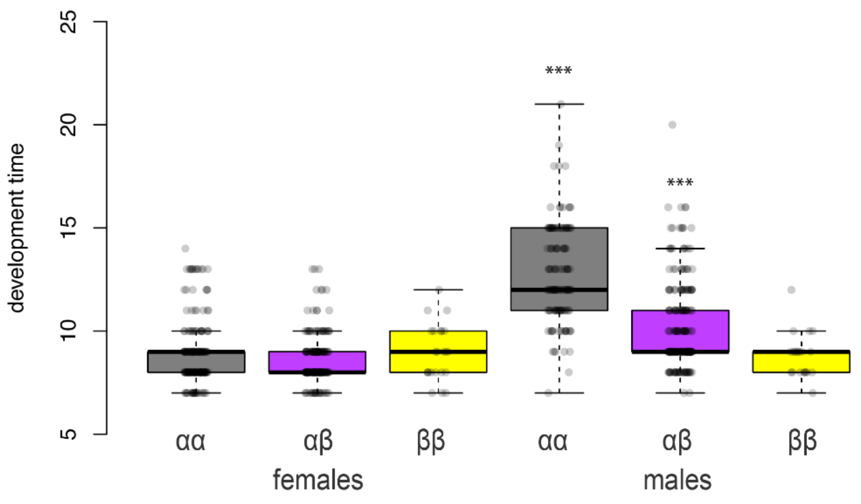

**Figure S3: Development time by genotype and sex.**

Time is counted as the number of days between egg laying and adult emergence. Data are drawn from the 5th generation, with the 16 replicates pooled. *Coelopa frigida* were raised on either Fucaceae or Laminariaceae, at low densities and constant temperature 25°C. Development time is expected to be longer under a higher density or at lower temperature. For instance in a similar experiment at 15°C, the females and the first males generally emerged after around 15 days while the last males took more than 30 days.

**Table S5: Analysis of development time**

**Generalized linear mixed model testing the effect of sex, genotype and substrate on development time.**

Box identity (replicate of the experimental evolution) was taken as random factor and we applied a Poisson transformation. $X^2$ and p-values come from comparisons between nested models with $X^2$-tests.

| | variable | DF | $X^2$ | P-value | |
|---|---|---|---|---|---|
| All samples | sex | 3 | 100.3 | <0.001 | |
| | genotype | 4 | 26.2 | <0.001 | |
| | substrat | 3 | 0.05 | 0.81 | |
| | sex* genotype | 7 | 157.37 | <0.001 | (best model) |
| Females | genotype | 4 | 1.1 | 0.57 | |
| | substrat | 3 | 0.04 | 0.85 | |
| | genotype * substrat | 7 | 0.72 | 0.87 | |
| Males | genotype | 4 | 55.7 | <0.001 | (best model) |
| | substrat | 3 | 0.19 | 0.66 | |
| | genotype * substrat | 7 | 1.25 | 0.74 | |

#### Deviation of genotypic proportion in the eggs in the experimental evolution

**Table S6: Analysis of the deviation of genotypic proportion in the eggs relatively to random expectations Linear mixed models testing the effect of genotype, sex, substrate or population on genotypic frequencies in the eggs relatively to Hardy-Weinberg proportions of the previous generation**

Random factor is the identity of the replicate box. Note: The effect of the interaction between genotype and generation is driven by the lower excess of αα at generation 3-4-5 compared to generation 1-2 (Fig. S4). However, such effect of generation should be interpreted with much caution since sample size is 4 replicates par generations and deviations at generation 3 to 5 are not as reliable as at earlier generation given the low absolute numbers of BB found in the eggs or the adults.

| | Df | F | P |
|---|---|---|---|
| genotype | 2 | 24.9 | <0.001 |
| generation | 1 | 10.5 | 0.002 |
| population | 1 | 1.93 | 0.17 |
| substrate | 1 | 0.03 | 0.87 |
| genotype * generation | 2 | 8.34 | <0.001 |
| genotype * population | 2 | 0.07 | 0.92 |
| genotype * substrate | 2 | 0.08 | 0.92 |

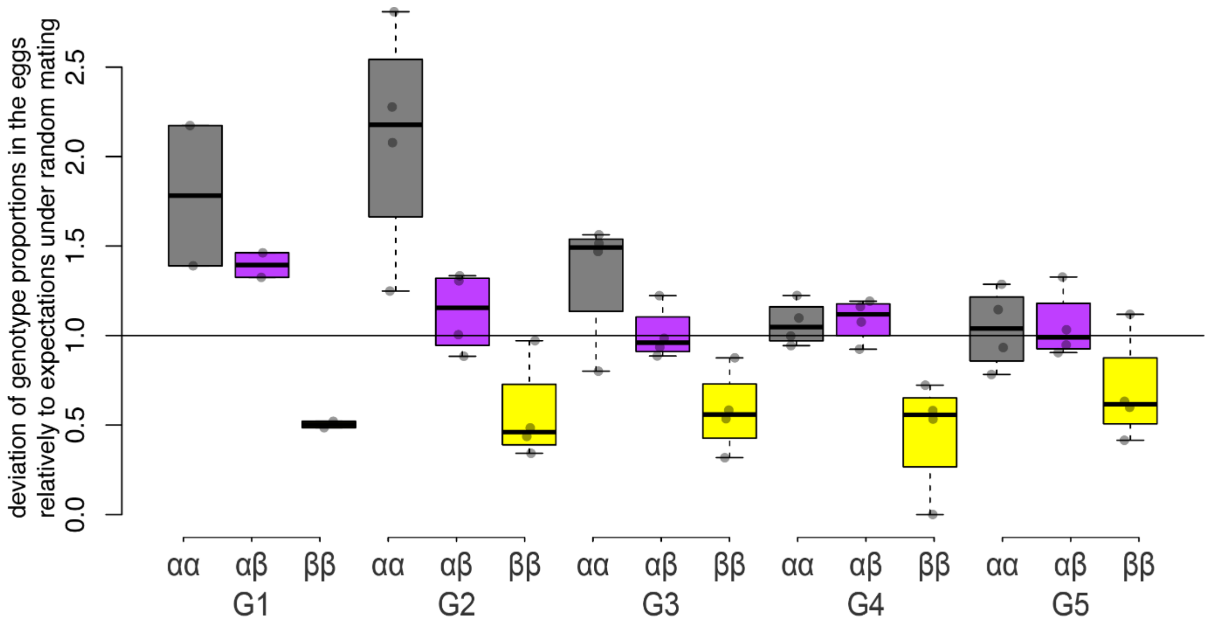

**Figure S4: Relative deviation from random expectations in the eggs for each genotype and each generation**

#### Modelling the evolution of inversion frequencies

**Table S7: Parameters of the individual-based model**

| | Parameter name | Description | Empirical values *[explored values]* used for simulations of the experiment | Default values [explored values] used for simulations in wild populations | theoretical models |
|---|---|---|---|---|---|
| All_models | G | Number of generations | 5 | 200 | 500 |
| | K | Number of eggs kept per generation | 1,000 | 10,000 | 10,000 |
| | E | Number of eggs per female | 70 | 70 | 70 |
| | V | Global egg-to-adult Viability | 0.3 | 0.3 | 0.3 |
| | R | Sex-ratio | 0.5 | 0.5 | 0.5 |
| | N0 | Number of adults at generation 0 | 300 | 3,000 | 3,000 |
| | P0-αα-ad | Proportions of αα adults at generation 0 | 0.07 | 0.07 | 0.25 |
| | P0-αβ-ad | Proportions of αβ adults at generation 1 | 0.49 | 0.49 | 0.5 |
| | P0-ββ-ad | Proportions of ββ adults at generation 2 | 0.44 | 0.44 | 0.25 |
| | Sααf | Relative egg-to-adult survival of αα females | 0.71 | 0.71 *[0.71; 0.52; 0.56]* | $1-s_f$ |
| | Sαβf | Relative egg-to-adult survival of αβ females | 0.9 | 0.90 *[0.90; 1.0; 1.0]* | $1-s_f*H_s$ |
| | Sββf | Relative egg-to-adult survival of ββ females | 1.0 | 1 *[1.0; 0.63; 0.34]* | 1 |
| | Sααm | Relative egg-to-adult survival of αα males | 0.81 | 0.81 *[0.81; 0.40; 0.33]* | $1-s_m$ |
| | Sαβm | Relative egg-to-adult survival of αβ males | 1.0 | 1.0 *[1.0; 1.0; 1.0]* | $1-s_m*H_s$ |
| | Sββm | Relative egg-to-adult survival of ββ males | 0.88 | 0.88 *[0.88; 0.35; 0.32]* | 1 |
| | Tααf | Relative fecundity of αα females | 1.0 | 1.0 | 1 |
| | Tαβf | Relative fecundity of αβ females | 0.97 | 0.97 | $1-t_f*H_t$ |
| | Tββf | Relative fecundity of ββ females | 0.87 | 0.87 | $1-t_f$ |
| | Tααm | Relative reproductive success of αα males | 1.0 *[1.0]* | 1.0 *[1.0]* | 1 |
| | Tαβm | Relative reproductive success of αβ males | *[0.525 : 1.0]* | 0.5 *[0.55 : 1.0]* | $1-t_m*H_t$ |
| | Tββm | Relative reproductive success of ββ males | *[0.05 : 1.0]* | 0.1 *[0.1 : 1.0]* | $1-t_m$ |
| with_freq-dpdce effect | FDC | Frequency-dependance coefficient | *[0 : 0.9]* | 0.9 | 0 |
| | tααm | Relative reproductive success of αα males | 1.0 *[1.0]* | 1.0 | |
| | tαβm | Relative reproductive success of αβ males | *[0.55 : 1.0]* | 0.8 | |
| | tββm | Relative reproductive success of ββ males | *[0.1 : 1.0]* | 0.6 | |
| with environmental effect | Df | Development time of females (in days) | 8.8 | 8.8 | 8.8 |
| | Dααm | Development time of αα males (in days) | 12.8 | 12.8 | 12.8 |
| | Dαβm | Development time of αβ males (in days) | 10.3 | 10.3 | 10.3 |
| | Dββm | Development time of ββ males (in days) | 8.7 | 8.7 | 8.7 |
| | DCV | Variation coefficient of development time | 0.5 | 0.5 | 0.5 |
| | Amean | Duration of habitat availability (in days) | 30 | 30 *[7 : 20]* | 30 |
| | ACV | Variation coefficient of the duration of habitat availability | 1 | 2 *[0 : 10]* | 1 |
| for theoretical model | $s / s_f / s_m$ | survival difference between homozygotes (for both sexes, females or males) | | | [0 : 1] |
| | $t / t_f / t_m$ | reproductive difference between homozygotes (for both sexes, females or males) | | | [0 : 1] |
| | $H_s$ | dominance in heterozygotes for survival | | | [-0,25, 0, 0.25, 0.5] |
| | $H_t$ | dominance in heterozygotes for reproduction | | | [-0,25, 0, 0.25, 0.5] |

**Table S8: Goodness of fit to empirical data between various scenarios on 5 generations.**
Fit is measured by the nrmse (normalized mean squared error) on the genotypic proportions in the adults and eggs. The difference of mean nrmse between all scenarios by a pairwise t-test between all the scenario (30 replicates per scenario), corrected for multiple testing following Benjamini & Hochberg. Below we present only the comparison to the best 6 models, selected as best based on their goodness of fit and different parametrizations explored. *** stands for p<0.01

| | | Scenario | $T_{\alpha\alpha\text{-}m}$ | $T_{\alpha\beta\text{-}m}$ | $T_{\beta\beta\text{-}m}$ | *FDC* | mean nrmse | p-value of t-test on the mean nrmse difference between each scenario the selected best 6 scenarios | | | | | |
|---|---|---|---|---|---|---|---|---|---|---|---|---|---|
| | | | | | | | | Exp BB0.1 | Exp BB0.05 | Exp Domin BB0.05 | Exp BB0.4 freq0.9 | Exp BB0.4 freq0.8 | Exp BB0.6 freq0.9 |
| ***BEST SCENARIOS FITTING EMPIRICAL OBSERVATIONS*** | | | | | | | | | | | | | |
| **Without frequency-dependance** | **Co-dominance** | exp_BB0.1 | 1 | 0.55 | 0.1 | 0 | **_28_** | | 0.53 | 0.34 | 0.50 | 0.41 | 0.78 |
| | | exp_BB0.05 | 1 | 0.525 | 0.05 | 0 | **_28_** | 0.53 | | 0.10 | 0.97 | 0.86 | 0.35 |
| | **dominance** | exp_domin_BB0.05 | 1 | 1 | 0.05 | 0 | **_33_** | 0.34 | 0.10 | | 0.10 | 0.07 | 0.51 |
| **With strong frequency-dependance effect** | | exp_BB0.4_freq0.9 | 1 | 0.7 | 0.4 | 0.9 | **_26_** | 0.50 | 0.97 | 0.10 | | 0.89 | 0.34 |
| | | exp_BB0.4_freq0.8 | 1 | 0.7 | 0.4 | 0.8 | **_26_** | 0.41 | 0.86 | 0.07 | 0.89 | | 0.27 |
| | | exp_BB0.6_freq0.9 | 1 | 0.8 | 0.6 | 0.9 | **_30_** | 0.78 | 0.35 | 0.51 | 0.34 | 0.27 | |
| ***OTHER SCENARIOS WITH REDUNDANT PARAMETRIZATION AND EQUIVALENT FIT*** | | | | | | | | | | | | | |
| **Equivalent to the case BB=0.1 and no frequency-dependence (very low BB success)** | | exp_BB0.1_freq0.2 | 1 | 0.55 | 0.1 | 0.2 | **_25_** | 0.21 | 0.55 | **0.02** | 0.58 | 0.68 | 0.12 |
| | | exp_BB0.1_freq0.3 | 1 | 0.55 | 0.1 | 0.3 | **_25_** | 0.29 | 0.68 | **0.04** | 0.71 | 0.82 | 0.17 |
| | | exp_BB0.1_freq0.4 | 1 | 0.55 | 0.1 | 0.4 | **_28_** | 0.52 | 1.00 | 0.10 | 0.98 | 0.87 | 0.35 |
| | | exp_BB0.1_freq0.5 | 1 | 0.55 | 0.1 | 0.5 | **_26_** | 0.35 | 0.77 | 0.05 | 0.80 | 0.91 | 0.22 |
| | | exp_BB0.1_freq0.6 | 1 | 0.55 | 0.1 | 0.6 | **_27_** | 0.64 | 0.87 | 0.15 | 0.85 | 0.74 | 0.45 |
| | | exp_BB0.1_freq0.1 | 1 | 0.55 | 0.1 | 0.1 | **_28_** | 0.93 | 0.59 | 0.29 | 0.57 | 0.47 | 0.71 |
| | | exp_BB0.1_freq0.7 | 1 | 0.55 | 0.1 | 0.7 | **_33_** | 0.25 | 0.07 | 0.85 | 0.06 | **0.04** | 0.39 |
| | | exp_BB0.2_freq0.6 | 1 | 0.6 | 0.2 | 0.6 | **_27_** | 0.46 | 0.92 | 0.08 | 0.95 | 0.94 | 0.30 |
| | | exp_BB0.2_freq0.5 | 1 | 0.6 | 0.2 | 0.5 | **_28_** | 0.97 | 0.55 | 0.32 | 0.53 | 0.43 | 0.76 |
| | | exp_BB0.2_freq0.8 | 1 | 0.6 | 0.2 | 0.8 | **_28_** | 0.71 | 0.81 | 0.18 | 0.78 | 0.67 | 0.51 |
| | | exp_BB0.2_freq0.7 | 1 | 0.6 | 0.2 | 0.7 | **_29_** | 0.85 | 0.40 | 0.45 | 0.38 | 0.31 | 0.93 |
| | | exp_BB0.2_freq0.4 | 1 | 0.6 | 0.2 | 0.4 | **_30_** | 0.64 | 0.26 | 0.64 | 0.25 | 0.19 | 0.85 |
| | | exp_BB0.2_freq0.9 | 1 | 0.6 | 0.2 | 0.9 | **_31_** | 0.71 | 0.31 | 0.57 | 0.29 | 0.23 | 0.93 |
| | | exp_BB0.2_freq0.3 | 1 | 0.6 | 0.2 | 0.3 | **_31_** | 0.61 | 0.24 | 0.67 | 0.23 | 0.18 | 0.82 |
| | | exp_BB0.2_freq0.2 | 1 | 0.6 | 0.2 | 0.2 | **_33_** | 0.20 | **0.05** | 0.75 | **0.05** | **0.03** | 0.32 |
| **Equivalent to the case with strong-frequency dependence effect (and higher BB success)** | | exp_BB0.6_freq0.8 | 1 | 0.8 | 0.6 | 0.8 | **_34_** | 0.10 | **0.02** | 0.50 | **0.02** | **0.01** | 0.17 |
| | | exp_BB0.4_freq0.7 | 1 | 0.7 | 0.4 | 0.7 | **_34_** | 0.17 | **0.04** | 0.69 | **0.04** | **0.03** | 0.28 |
| | | exp_BB0.8_freq0.9 | 1 | 0.9 | 0.8 | 0.9 | **_34_** | 0.20 | 0.05 | 0.76 | **0.05** | **0.03** | 0.33 |
| ***OTHER SCENARIOS EXPLORED*** | | | | | | | | | | | | | |
| **Combination of parameters with significantly-reduced fit to empirical observations (by comparison with best models)** | | exp_BB0.1_freq0.8 | 1 | 0.55 | 0.1 | 0.8 | **_35_** | 0.09 | **0.02** | 0.49 | **0.02** | **0.01** | 0.17 |
| | | exp_BB0.2_freq0.1 | 1 | 0.6 | 0.2 | 0.1 | **_35_** | 0.07 | **0.01** | 0.43 | **0.01** | **0.01** | 0.14 |
| | | exp_BB0.4_freq0.6 | 1 | 0.7 | 0.4 | 0.6 | **_37_** | **0.01** | *** | 0.10 | *** | *** | **0.02** |
| | | exp_domin_BB0.1 | 1 | 1 | 0.1 | 0 | **_37_** | **0.01** | *** | 0.15 | *** | *** | **0.03** |
| | | exp_BB0.1_freq0.9 | 1 | 0.55 | 0.1 | 0.9 | **_38_** | **0.02** | *** | 0.15 | *** | *** | **0.03** |
| | | exp_BB0.2 | 1 | 0.6 | 0.2 | 0 | **_38_** | **0.01** | *** | 0.11 | *** | *** | **0.02** |
| | | exp_BB0.2_freq0.0 | 1 | 0.6 | 0.2 | 0 | **_39_** | *** | *** | **0.04** | *** | *** | **0.01** |
| | | exp_BB1.0_freq0.9 | 1 | 1 | 1 | 0.9 | **_40_** | *** | *** | **0.02** | *** | *** | *** |
| | | exp_BB0.6_freq0.7 | 1 | 0.8 | 0.6 | 0.7 | **_40_** | *** | *** | **0.01** | *** | *** | *** |

| | | | | | | | | | | |
|---|---|---|---|---|---|---|---|---|---|---|
| exp_BB0.4_freq0.5 | 1 | 0.7 | 0.4 | 0.5 | ***41*** | *** | *** | 0.01 | *** | *** | *** |
| exp_BB0.8_freq0.8 | 1 | 0.9 | 0.8 | 0.8 | ***44*** | *** | *** | *** | *** | *** | *** |
| exp_domin_BB0.2 | 1 | 1 | 0.2 | 0 | ***44*** | *** | *** | *** | *** | *** | *** |
| exp_BB0.4_freq0.4 | 1 | 0.7 | 0.4 | 0.4 | ***45*** | *** | *** | *** | *** | *** | *** |
| exp_BB0.4_freq0.3 | 1 | 0.7 | 0.4 | 0.3 | ***48*** | *** | *** | *** | *** | *** | *** |
| exp_BB1.0_freq0.8 | 1 | 1 | 1 | 0.8 | ***49*** | *** | *** | *** | *** | *** | *** |
| exp_BB0.8_freq0.7 | 1 | 0.9 | 0.8 | 0.7 | ***50*** | *** | *** | *** | *** | *** | *** |
| exp_BB0.3 | 1 | 0.65 | 0.3 | 0 | ***51*** | *** | *** | *** | *** | *** | *** |
| exp_BB0.6_freq0.6 | 1 | 0.8 | 0.6 | 0.6 | ***51*** | *** | *** | *** | *** | *** | *** |
| exp_BB0.6_freq0.5 | 1 | 0.8 | 0.6 | 0.5 | ***52*** | *** | *** | *** | *** | *** | *** |
| exp_domin_BB0.3 | 1 | 1 | 0.3 | 0 | ***53*** | *** | *** | *** | *** | *** | *** |
| exp_BB0.4_freq0.2 | 1 | 0.7 | 0.4 | 0.2 | ***53*** | *** | *** | *** | *** | *** | *** |
| exp_BB0.4_freq0.1 | 1 | 0.7 | 0.4 | 0.1 | ***55*** | *** | *** | *** | *** | *** | *** |
| exp_BB0.8_freq0.6 | 1 | 0.9 | 0.8 | 0.6 | ***58*** | *** | *** | *** | *** | *** | *** |
| exp_domin_BB0.4 | 1 | 1 | 0.4 | 0 | ***60*** | *** | *** | *** | *** | *** | *** |
| exp_BB1.0_freq0.7 | 1 | 1 | 1 | 0.7 | ***60*** | *** | *** | *** | *** | *** | *** |
| exp_BB0.6_freq0.4 | 1 | 0.8 | 0.6 | 0.4 | ***60*** | *** | *** | *** | *** | *** | *** |
| exp_BB0.4_freq0.0 | 1 | 0.7 | 0.4 | 0 | ***62*** | *** | *** | *** | *** | *** | *** |
| exp_BB0.4 | 1 | 0.7 | 0.4 | 0 | ***63*** | *** | *** | *** | *** | *** | *** |
| exp_BB0.8_freq0.5 | 1 | 0.9 | 0.8 | 0.5 | ***65*** | *** | *** | *** | *** | *** | *** |
| exp_BB0.6_freq0.3 | 1 | 0.8 | 0.6 | 0.3 | ***67*** | *** | *** | *** | *** | *** | *** |
| exp_domin_BB0.5 | 1 | 1 | 0.5 | 0 | ***69*** | *** | *** | *** | *** | *** | *** |
| exp_domin_BB0.6 | 1 | 1 | 0.6 | 0 | ***70*** | *** | *** | *** | *** | *** | *** |
| exp_BB0.6_freq0.2 | 1 | 0.8 | 0.6 | 0.2 | ***70*** | *** | *** | *** | *** | *** | *** |
| exp_BB1.0_freq0.6 | 1 | 1 | 1 | 0.6 | ***70*** | *** | *** | *** | *** | *** | *** |
| exp_BB0.5 | 1 | 0.75 | 0.5 | 0 | ***73*** | *** | *** | *** | *** | *** | *** |
| exp_BB0.8_freq0.4 | 1 | 0.9 | 0.8 | 0.4 | ***74*** | *** | *** | *** | *** | *** | *** |
| exp_BB1.0_freq0.5 | 1 | 1 | 1 | 0.5 | ***79*** | *** | *** | *** | *** | *** | *** |
| exp_BB0.6_freq0.1 | 1 | 0.8 | 0.6 | 0.1 | ***83*** | *** | *** | *** | *** | *** | *** |
| exp_BB0.8_freq0.3 | 1 | 0.9 | 0.8 | 0.3 | ***84*** | *** | *** | *** | *** | *** | *** |
| exp_BB0.6 | 1 | 0.8 | 0.6 | 0 | ***85*** | *** | *** | *** | *** | *** | *** |
| exp_domin_BB0.7 | 1 | 1 | 0.7 | 0 | ***86*** | *** | *** | *** | *** | *** | *** |
| exp_BB0.6_freq0.0 | 1 | 0.8 | 0.6 | 0 | ***86*** | *** | *** | *** | *** | *** | *** |
| exp_BB0.8_freq0.2 | 1 | 0.9 | 0.8 | 0.2 | ***86*** | *** | *** | *** | *** | *** | *** |
| exp_BB1.0_freq0.4 | 1 | 1 | 1 | 0.4 | ***87*** | *** | *** | *** | *** | *** | *** |
| exp_BB1.0_freq0.3 | 1 | 1 | 1 | 0.3 | ***89*** | *** | *** | *** | *** | *** | *** |
| exp_BB0.8_freq0.1 | 1 | 0.9 | 0.8 | 0.1 | ***94*** | *** | *** | *** | *** | *** | *** |
| exp_domin_BB0.8 | 1 | 1 | 0.8 | 0 | ***95*** | *** | *** | *** | *** | *** | *** |
| exp_BB0.7 | 1 | 0.85 | 0.7 | 0 | ***96*** | *** | *** | *** | *** | *** | *** |
| exp_BB0.8 | 1 | 0.9 | 0.8 | 0 | ***99*** | *** | *** | *** | *** | *** | *** |
| exp_BB0.9 | 1 | 0.95 | 0.9 | 0 | ***99*** | *** | *** | *** | *** | *** | *** |
| exp_BB1.0_freq0.2 | 1 | 1 | 1 | 0.2 | ***101*** | *** | *** | *** | *** | *** | *** |
| exp_BB0.8_freq0.0 | 1 | 0.9 | 0.8 | 0 | ***102*** | *** | *** | *** | *** | *** | *** |
| exp_domin_BB0.9 | 1 | 1 | 0.9 | 0 | ***105*** | *** | *** | *** | *** | *** | *** |
| exp_BB1.0_freq0.1 | 1 | 1 | 1 | 0.1 | ***105*** | *** | *** | *** | *** | *** | *** |
| exp_BB1.0 | 1 | 1 | 1 | 0 | ***105*** | *** | *** | *** | *** | *** | *** |
| exp_domin_BB1.0 | 1 | 1 | 1 | 0 | ***111*** | *** | *** | *** | *** | *** | *** |
| exp_BB1.0_freq0.0 | 1 | 1 | 1 | 0 | ***117*** | *** | *** | *** | *** | *** | *** |

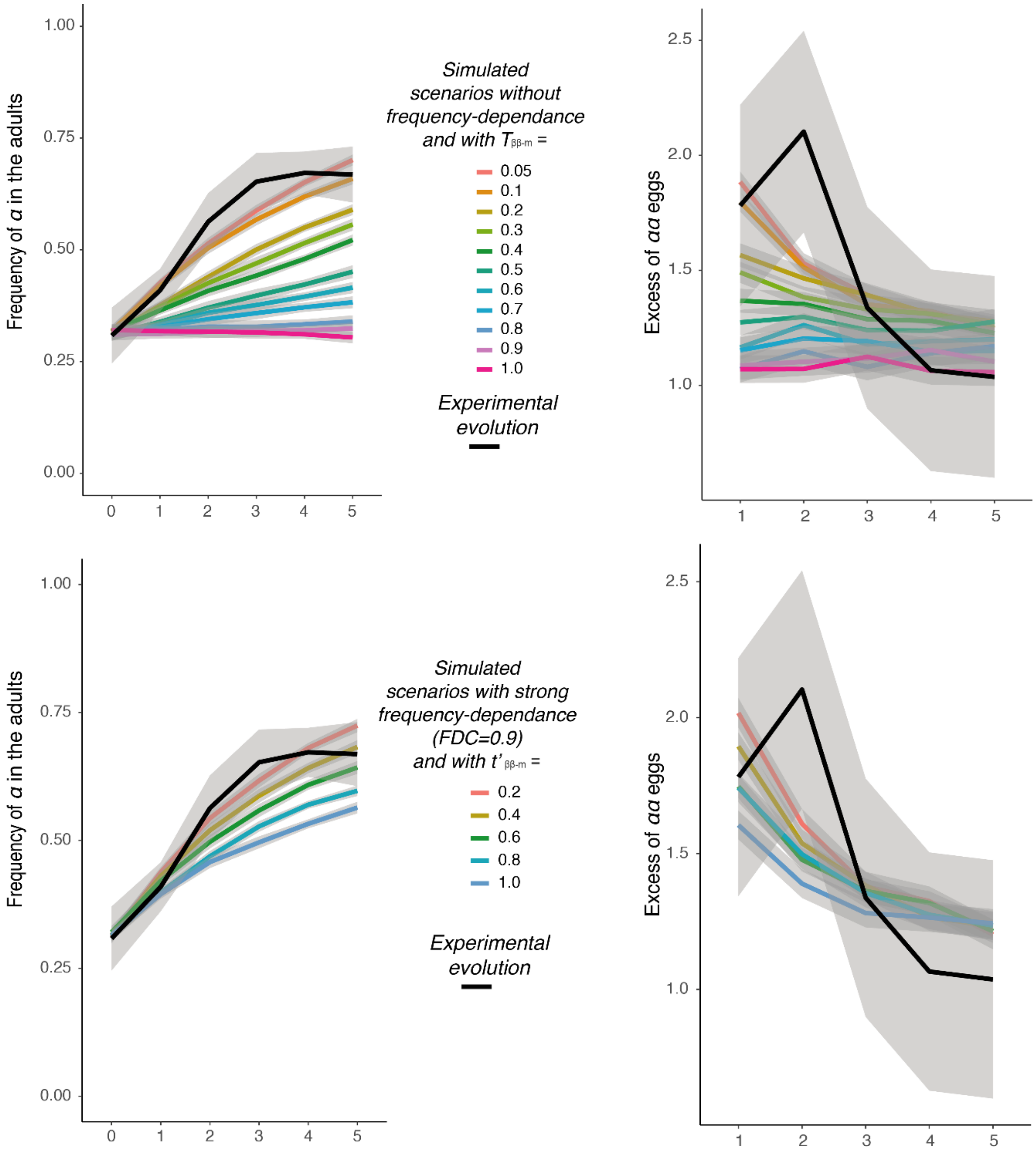

**Figure S5: Comparison of in silico experimental evolution to in vivo data**

Evolution of the frequency of the α rearrangement and the deviation of αα in the eggs across 5 generations. The two-best scenario (as scored by nRMSE) are TBB-m = 0.1 or 0.05 without frequency dependence, and FDC=0.9 and TBB-m = 0.4 or 0.6, with frequency-dependance.

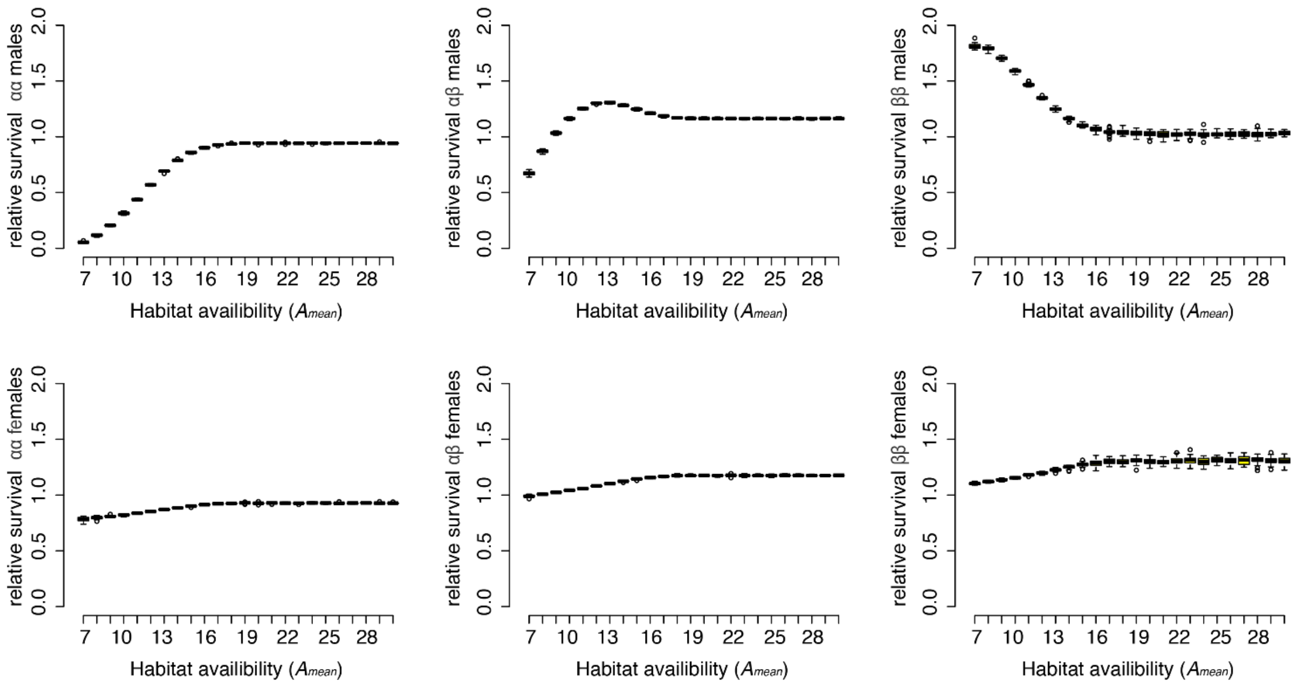

**Figure S6: Relative survival rate according to the duration of habitat availability.**

This value is an estimate of the relative survival rate includes both the intrinsic egg-to-adult survival rate and the effect of the environment. It is calculated as the deviation between the proportions in the adults and in the eggs.

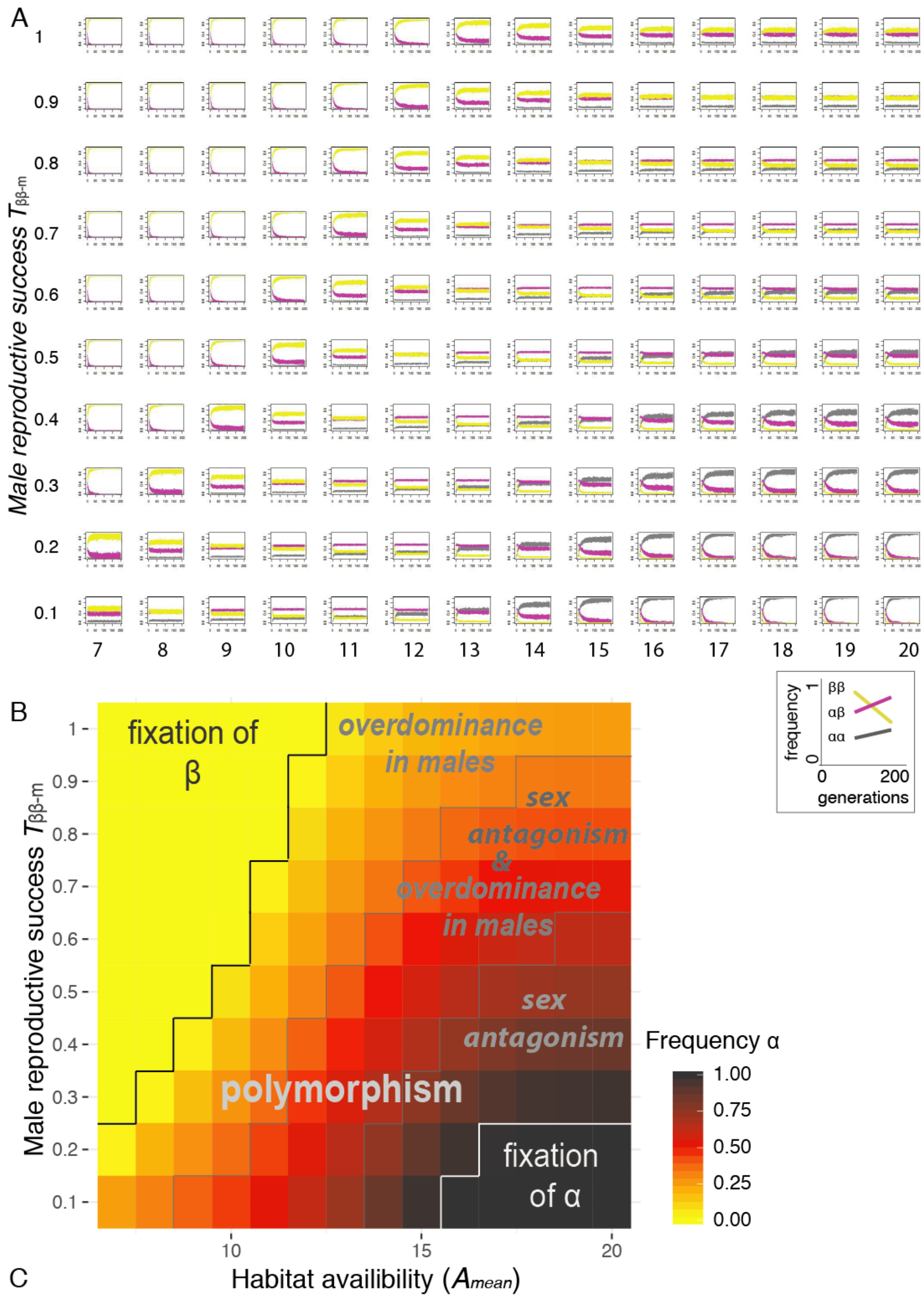

**Figure S7: Evolution of the three genotypes proportions in simulations co-varying male reproductive success and environment**.

Each subplot on panel (A) represents the evolution of the frequency of the three genotypes (ββ: yellow, αβ: purple, αα: grey) as a function of time (generations 0 to 200) as outline in the insert in the lowest left corner. The disposition of the plots is a mirror of Fig. 5A (recalled in panel B), that represent the frequency of α allele at equilibrium (after 200 generations). Each row represents a different value of Tββ-m (male reproductive success of ββ), from 1.0 (= Tαα-m) to 0.1 (= tenfold lower than Tαα-m). Each column represents a different value of the duration of habitat availability (A mean). Variability in the duration of the environment is fixed at Avar =2. Relative survival rate (S parameter) corresponds to values as estimated in the experiment (low density scenario)

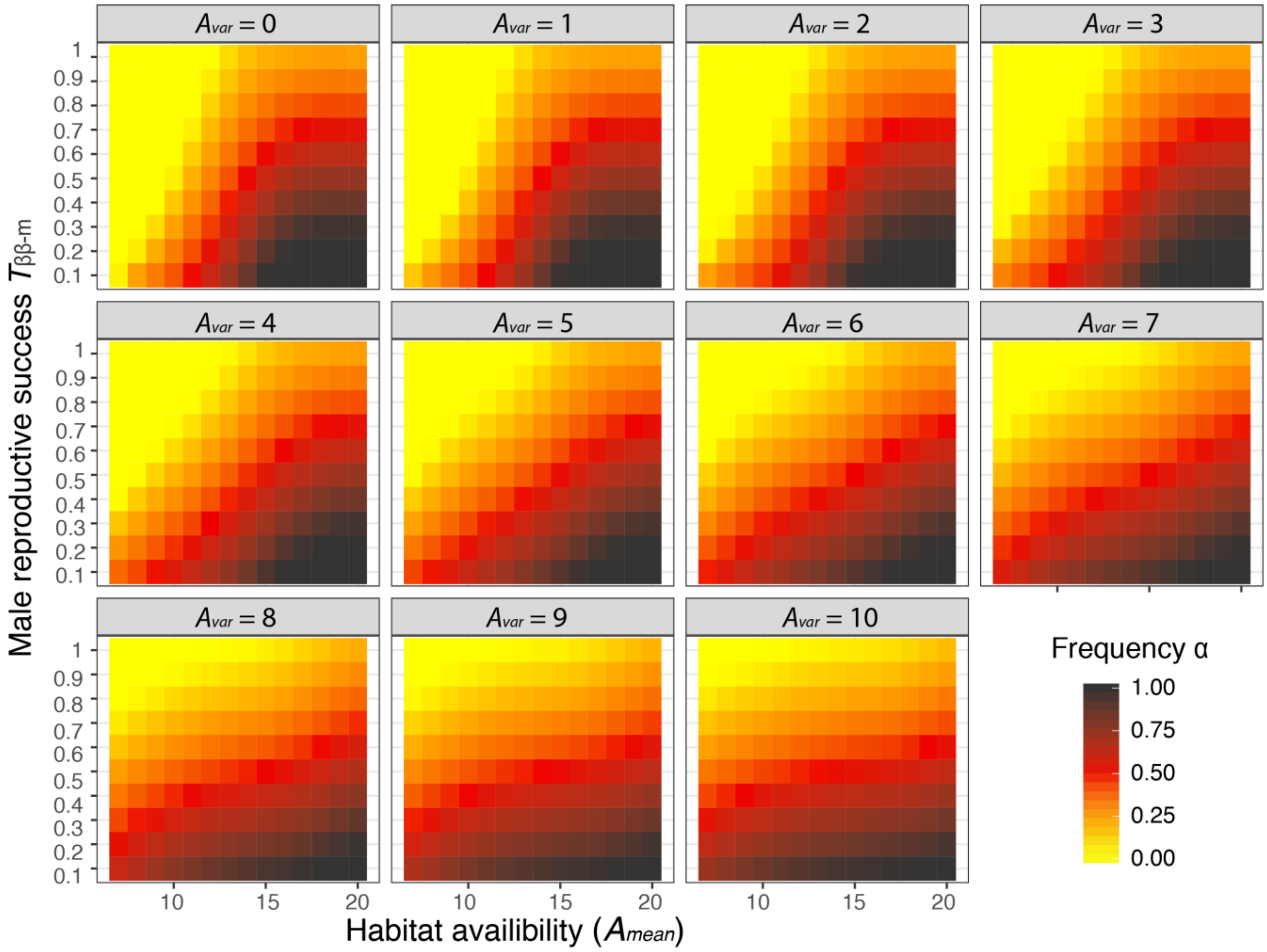

**Figure S8: Frequency of the α rearrangement in simulations varying male reproductive success, the mean duration of habitat availability (Amean) and its variability (Avar).**

Each subplot represent the mean frequency of alpha after 200 generations across 100 replicates in the parameter space defined by the y-axis, male reproductive success (Tββ-m), and the x-axis, mean duration of habitat availability (Amean). Yellow areas correspond to a fixation of the β allele, black area to the fixation of α allele, and orang/red/brown area to the persistence of polymorphism, as shown on Fig. 5A. With increasing variability in the duration of habitat availability (Avar, expressed in days), we observed a wider set of parameters for which polymorphism is maintained and α frequency remains at intermediate values.

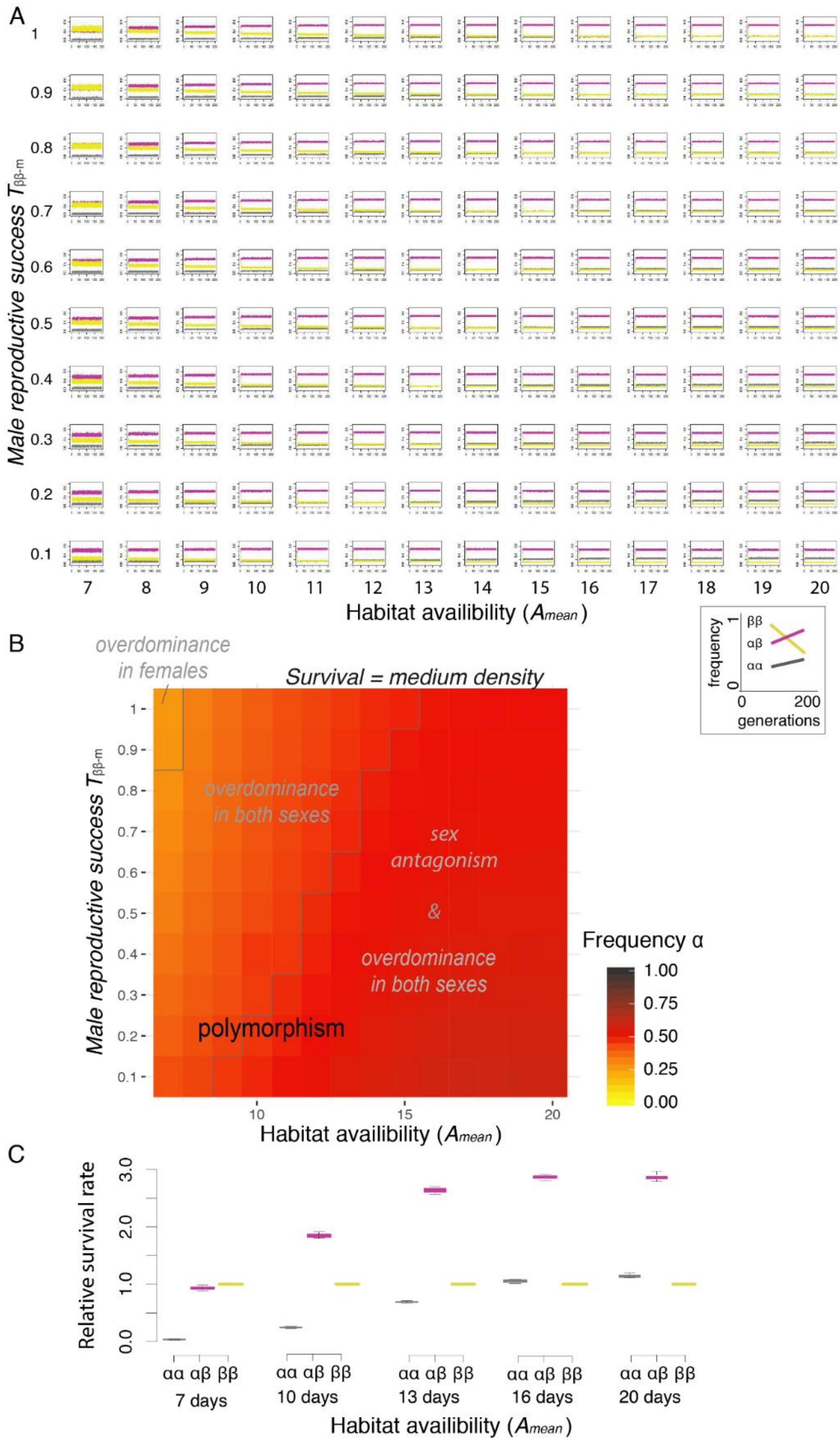

**Figure S9: Evolution of the three genotypes proportions in simulations co-varying male reproductive success and environment in the medium density scenario.**

Each subplot on panel (A) represents the evolution of the frequency of the three genotypes (ββ: yellow, αβ: purple, αα: grey) as a function of time (generations 0 to 200) as outline in the insert in the lowest left corner. The disposition of the plots is a mirror of panel B that represent the frequency of α allele at equilibrium (after 200 generations). Variability in the duration of the environment is fixed at Avar =2. Relative survival rate (S parameter) corresponds to values estimates at medium density by Butlin et al (10). This relative survival rate is further affected by the possibility to reach adulthood, as modelled with the interaction between development time and the availability of the habitat. Panel C thus represent the Overall relative survival rate in males, after the effect of environment, for the three genotypes in those simulations varying the duration of habitat availability (normalized relatively to Sββ-m = 1).

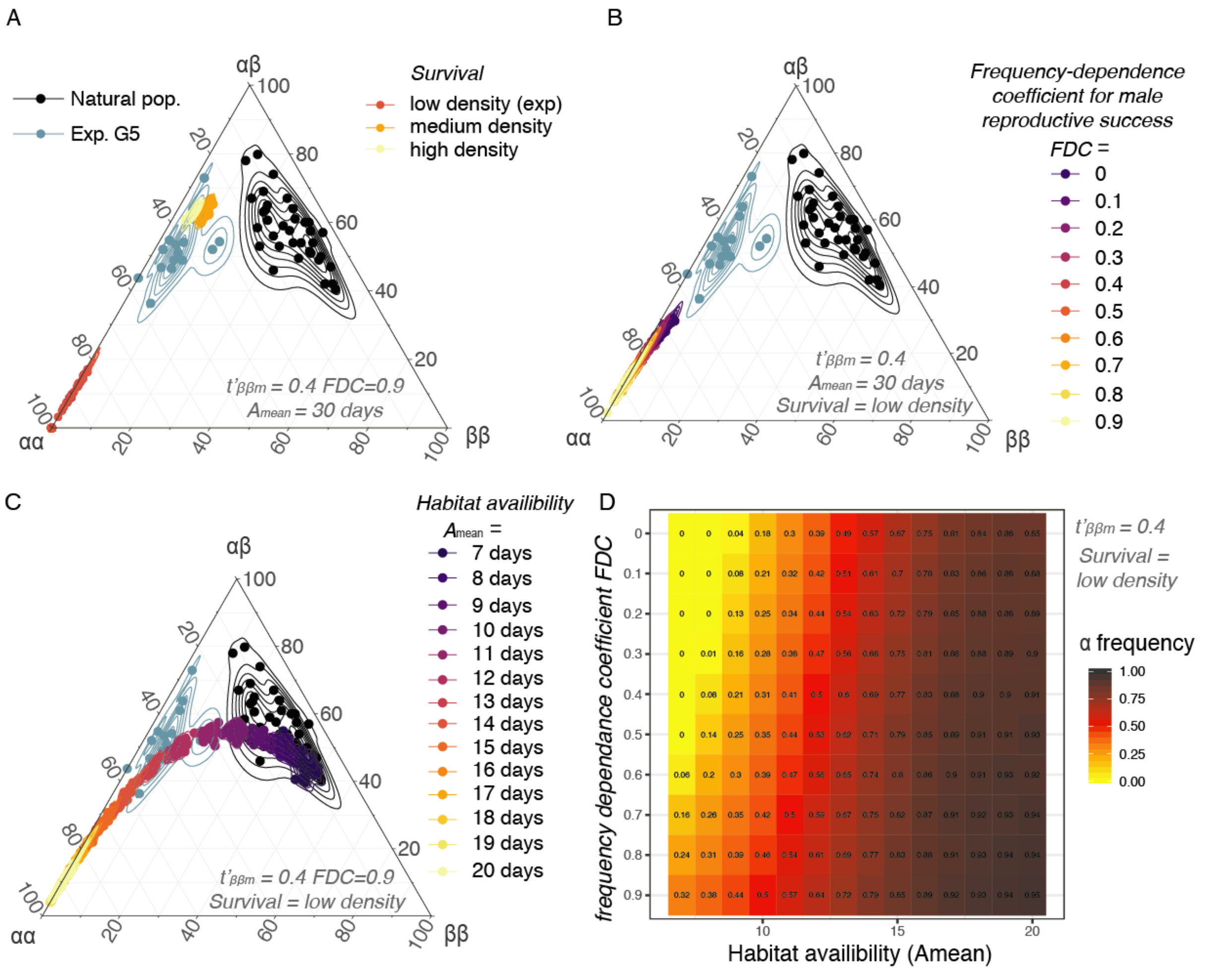

**Figure S10: Outcome of simulations taking into account frequency-dependence effect on male reproductive success**

(A-C)Ternary plots comparing the proportions of the three genotypes in natural populations (1, 11), after the 5th generation of our laboratory experiment and at the equilibrium after 200 generations of simulations taking into account a effect of the frequency of αα males on male reproductive success. In a few words, the relative reproductive success of αα (large) males is reduced when they are more frequent. Since this parameter is fixed to 1, this translate into the reproductive success of ββ males increasing with the frequency of αα males. The set of parameters is inspired by the best model fitting experimental data (see Table S8 & Fig S5), i. e. FDC=0.9 and t'ββ-m =0.4. We explore scenarios varying (A) the effect of density, and the related relative survival rate, (B) the range of values for the frequency-dependence parameter on male relative reproductive success (FDc, t'ββ-m is fixed to 0.4) C) the effect of a limited duration of the habitat availability (Amean = [7-20 days], Avar = 2 days). (D) Polymorphism persistence and frequency of the inversion at equilibrium in simulations co-varying the duration of habitat availability (which modulate male relative survival) and the strength of the frequency-dependence impact on male reproductive success.

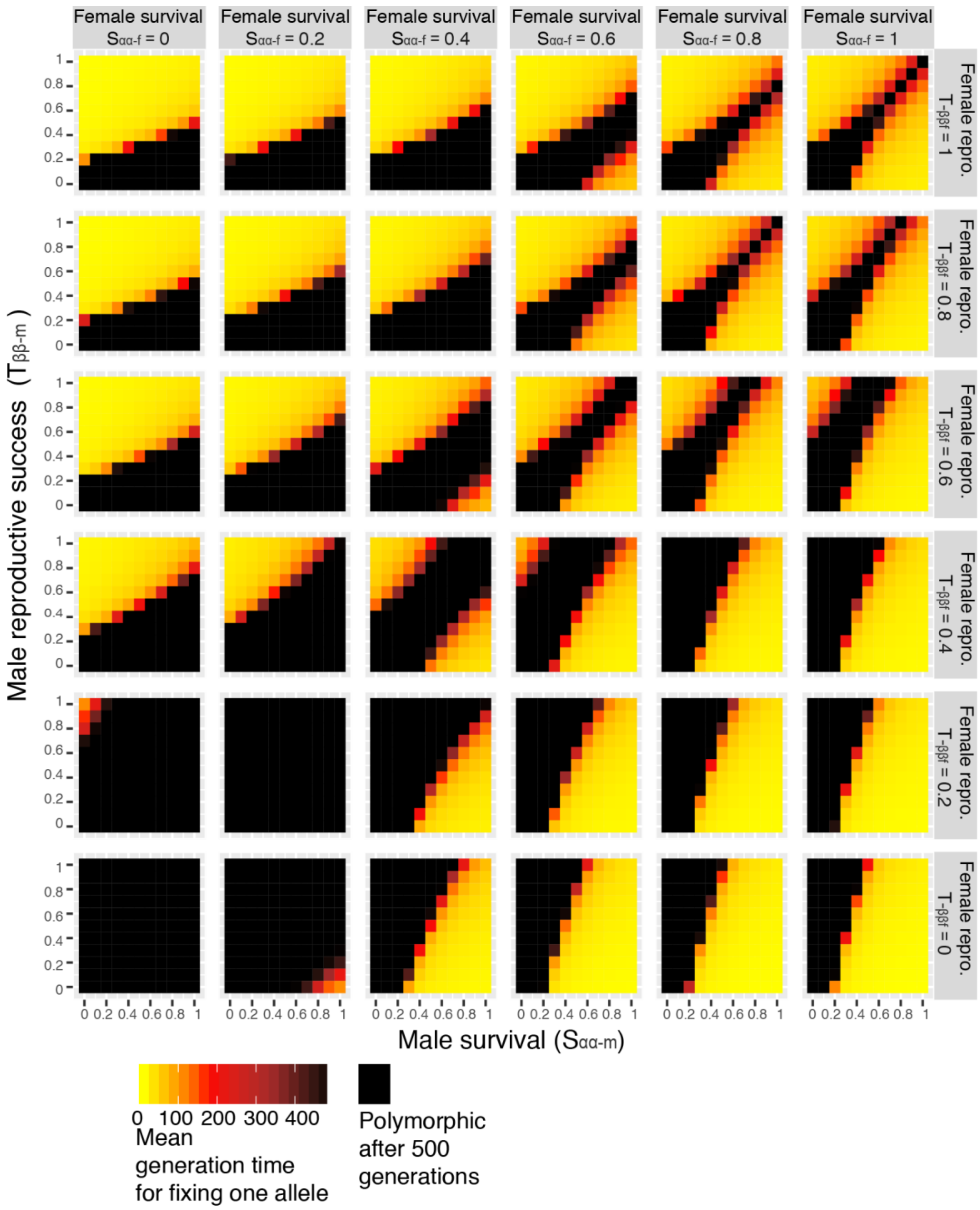

**Figure S11: Time to fixation with sexually-varying fitness parameters**
